# Modeling human *TBX5* haploinsufficiency predicts regulatory networks for congenital heart disease

**DOI:** 10.1101/835603

**Authors:** Irfan S. Kathiriya, Kavitha S. Rao, Giovanni Iacono, W. Patrick Devine, Andrew P. Blair, Swetansu K. Hota, Michael H. Lai, Bayardo I. Garay, Reuben Thomas, Henry Z. Gong, Lauren K. Wasson, Piyush Goyal, Tatyana Sukonnik, Gunes A. Akgun, Laure D. Bernard, Brynn N. Akerberg, Fei Gu, Kai Li, William T. Pu, Joshua M. Stuart, Christine E. Seidman, J. G. Seidman, Holger Heyn, Benoit G. Bruneau

## Abstract

Haploinsufficiency of transcriptional regulators causes human congenital heart disease (CHD). However, underlying CHD gene regulatory network (GRN) imbalances are unknown. Here, we define transcriptional consequences of reduced dosage of the CHD-linked transcription factor, TBX5, in individual cells during cardiomyocyte differentiation from human induced pluripotent stem cells (iPSCs). We discovered highly sensitive dysregulation of TBX5-dependent pathways— including lineage decisions and genes associated with cardiomyocyte function and CHD genetics—in discrete subpopulations of cardiomyocytes. GRN analysis identified vulnerable nodes enriched for CHD genes, indicating that cardiac network stability is sensitive to TBX5 dosage. A GRN-predicted genetic interaction between *Tbx5* and *Mef2c* was validated in mouse, manifesting as ventricular septation defects. These results demonstrate exquisite sensitivity to TBX5 dosage by diverse transcriptional responses in heterogeneous subsets of iPSC-derived cardiomyocytes. This predicts candidate GRNs for human CHDs, with implications for quantitative transcriptional regulation in disease.

## Introduction

CHDs are a leading cause of childhood morbidity and mortality, and incidence of CHDs is estimated to be ten-fold higher in human fetuses of spontaneous termination (Hoffman, 1995; Hoffman and Kaplan, 2002). Many human mutations linked to CHDs are predicted to result in reduced dosage of transcriptional regulators, including transcription factors (TFs) and chromatin-modifying genes (Zaidi and Brueckner, 2017). Despite advances to elucidate the roles of individual factors in heart development and CHDs, how dosage of transcriptional regulators translates to altered GRNs is not known.

In humans, homozygous loss of function (LOF) mutations of *TBX5* are not observed in the genome aggregation database of exomes and genomes from large-scale sequencing projects (Karczewski et al., 2020) and is presumed to cause fetal demise. In conjunction, *Tbx5* null mice die of embryonic lethality from severely deformed hearts (Bruneau et al., 2001). Heterozygous mutations in the T-box TF gene *TBX5* cause Holt-Oram syndrome (HOS) (Basson et al., 1997; Li et al., 1997), which uniformly presents with upper limb defects and often with CHDs that include ventricular or atrial septal defects, diastolic dysfunction and arrhythmias. Experiments in mice have revealed a stepwise sensitivity to reductions in Tbx5 dosage (Bruneau et al., 2001; Mori et al., 2006). These findings demonstrate that a reduction in TBX5 dosage perturbs downstream gene expression. However, the disrupted regulatory networks and mechanisms are not understood.

To build upon findings from mouse models, a tractable human model system is required to study human *TBX5* haploinsufficiency. Human heart tissue from normal, living individuals is largely inaccessible for molecular analysis. As the estimated prevalence of Holt-Oram syndrome is 1:100,000, pathological or surgical specimens from affected patients are very limited. Alternatively, genome editing in human induced pluripotent stem (iPS) cells enables targeted genetic manipulations in an isogenic background. Furthermore, these targeted mutant iPSCs can be differentiated into varied cardiac cell types, including cardiomyocytes, and then subjected to single cell RNA sequencing (RNA-seq). This *in vitro* system provides a promising human cellular platform for gene-centered cardiac disease modeling at single cell resolution. Although iPSC-derived cardiomyocyte differentiation lacks a three-dimensional context for patterning and organization of myriad cell types of the heart, it recapitulates key developmental steps, including mesendoderm and cardiac precursors. Directed cardiomyocyte differentiation leads to a predominance of ventricular cardiomyocytes, with production of some atrial cardiomyocytes and surrounding cell types, such as fibroblasts, endothelial cells and endodermal cells, providing a useful multicellular system to model aspects of human cardiogenesis.

In considering how *TBX5* haploinsufficiency might cause CHDs, at least two scenarios are possible: 1. Reduced dosage may only affect genes in vulnerable cell types in specific anatomical locations, such as the ventricular chamber or septum. 2. Reduced dosage may affect cardiac gene expression broadly, but altered programs manifest as morphologic defects only in cell types of anatomic structures most sensitive to the disturbance. The first scenario would be challenging to investigate in two-dimensional cultures, particularly if susceptible region-specific cell types are absent. The second predicts that changes in gene expression might be detected by bulk RNA-seq studies of heterozygous human iPS cell models of CHDs (Theodoris et al., 2015; Ang et al., 2016; Gifford et al., 2019), but relevant, discrete alterations in a complex cell mixture could be missed. Discerning between these scenarios will require a single cell analysis approach.

Here, we used an allelic series of *TBX5* in engineered human iPS cells, comprising wildtype, and heterozygous or homozygous loss of function mutations, to investigate GRNs that are altered in response to reduced TBX5 dosage. We observed TBX5 dose-dependent cellular phenotypes reminiscent of anomalies in patients with *TBX5* mutations. We deployed single cell RNA-seq across a time course of differentiation and observed that the acquisition of ventricular cardiomyocyte fate is sensitive to TBX5 dosage. We also discovered discrete gene expression responses to reduced *TBX5* dosage in cardiomyocyte subpopulations. From these data, we identified putative cardiac GRNs that help explain several cellular phenotypes related to human CHD. We validated one of these GRN-predicted genetic interactions in mice. *Tbx5* and *Mef2c* interact to cause muscular ventricular septal defects (VSDs), a common type of human CHD. We conclude that TBX5 dosage sensitivity, modeled in human iPS cells, reveals discrete gene regulation programs in an unanticipated variety of cardiomyocyte subtypes and informs the biology of human CHD.

## Results

### Impaired human cardiomyocyte differentiation and function by reduced TBX5 dosage

To determine a role for TBX5 dosage in human cardiac biology, we created an isogenic allelic series of human iPS cells mutant for *TBX5,* using CRISPR/Cas9-mediated genome editing to target exon 3 of *TBX5* at the start of the essential T-box domain (Figure S1A, B). We isolated targeted iPS cell lines, including heterozygous (*TBX5 ^in/+^*) and homozygous (*TBX5^in/del^* and *TBX5^PuR/PuR^*) mutants (Figure 1A, S1C-F). We also isolated a control (*TBX5^+/+^*) iPS cell line, which was exposed to CRISPR/Cas9 nuclease but not mutated at exon 3 of *TBX5*, to control for off-target effects and the sub-cloning procedure. Subsequently, we refer to wildtype and control collectively as “WT” when significant differences between them were not observed. TBX5 protein levels in cardiomyocytes differentiated from these lines were diminished in *TBX5^in^*^/+^ cells and absent in *TBX5^in^*^/*del*^ and *TBX5^PuR/PuR^* cells (Figure 1B), consistent with a dosage-step allelic series of mutant *TBX5* loss-of-function cell lines.

**Figure 1.**
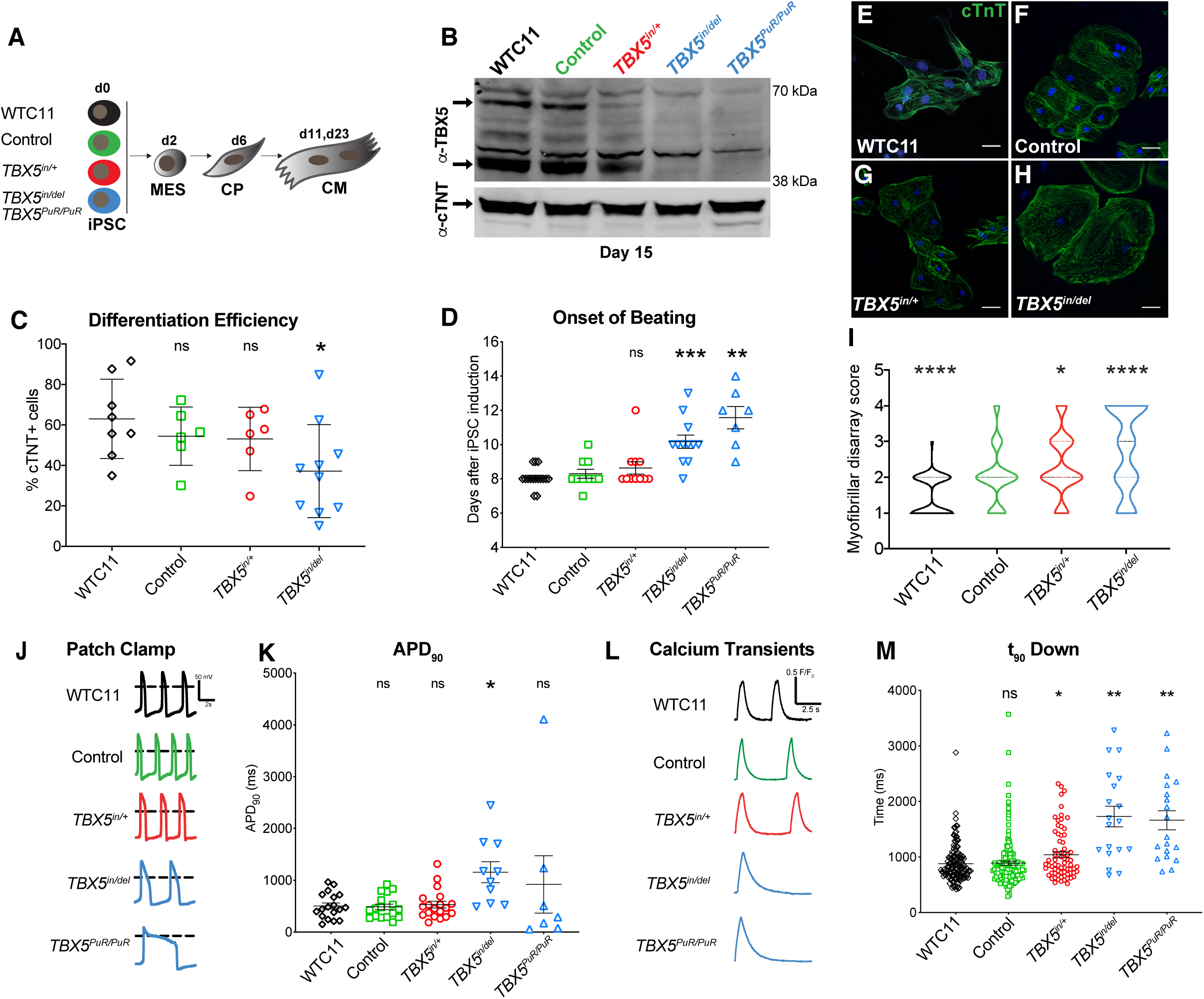
A human allelic series of *TBX5* mutants model features of congenital heart disease. (A) Parental iPS cell line WTC11, control (CRISPR-treated, unmodified at exon 3 of *TBX5*) and targeted *TBX5* loss of function mutants (*TBX5^in/+^*, *TBX5^in/del^*, or *TBX5^PuR/PuR^*) underwent directed differentiation to cardiomyocytes (CM) via mesoderm (MES) and cardiac precursor (CP) stages. (B) TBX5 and cTNT protein expression for each *TBX5* genotype from the cardiomyocyte stage at day 15. (C) Differentiation efficiency by flow cytometry for cTNT^+^ cells (* p-value<0.05 by unpaired *t* test). (D) Onset of beating (** p-value<0.01, *** p-value<0.001 by unpaired *t* test). (E-I) Myofibrillar arrangement of cardiomyocytes (* p-value<0.05, **** p-value<0.0001 by Fisher’s exact test). (J) Action potentials by patch clamp of single beating cells for each *TBX5* genotype. (K) Action potential duration at 90% (APD_90_) (* FDR<0.05). (L, M) Traces of calcium transients from single or small clusters of beating cells were analyzed, including time at 90% decay (t_90_ down) (* FDR<0.05, ** FDR<0.01). Error bars represent standard deviation (J, K) or standard error (L, M) of the mean. Data for *TBX5^PuR/PuR^* is shown where available.

We observed reduced cardiomyocyte differentiation efficiency and a delay in onset of beating by loss of *TBX5*, when compared to WT (Figure 1C, D). Worsening sarcomere disarray was seen by stepwise depletion of *TBX5* (Figure 1E-I). Patch clamp analysis of cardiomyocytes, which were predominantly ventricular in this differentiation method, revealed lengthened action potential duration (APD) in *TBX5^in^*^/*del*^ cells (Figure 1J, K; action potential duration at 90% repolarization (APD_90_), adj p-value<0.04 by Holm-Sidak test) (Holm1979), consistent with previous findings (Churko et al., 2018; Karakikes et al., 2017). Although *TBX5^PuR^*^/*PuR*^ cells showed high variability of APD_90_ and were not statistically significantly different from WT, some recordings were distinctly abnormal, displaying striking APD_90_ durations eight times greater than an average wildtype or control cell (Figure 1J, K). Calcium imaging of spontaneously beating cardiomyocytes revealed protracted calcium transient durations in *TBX5^in^*^/*del*^ and *TBX5^PuR^*^/*PuR*^ cells (time of 90% decay; (t_90_ down), adj p-value<9E-4 by Holm-Sidak test), with an intermediate defect in *TBX5^in/+^* cells (adj p-value<0.01) (Figure 1L, M), implying a potential impairment of cardiomyocyte relaxation. Together, these cellular findings recapitulated several pathological characteristics, which may contribute to diastolic dysfunction in HOS in mice and humans (Zhou et al., 2005; Zhu et al., 2008).

### Resolving susceptible cardiac cell types from reduced TBX5 dosage

To determine how TBX5 dosage alters gene expression in a heterogeneous cell population, we used a droplet-based single cell RNA-seq method with cells collected from parental WTC11, control *TBX5*^+/+^, and mutant *TBX5* (*TBX5^in/+^*, *TBX5^in^*^/*del*^) genotypes. From three time points during cardiomyocyte differentiation, we interrogated 55,782 cells with an average normalized read depth of 88,234 reads per cell (Figure 2A-C). At day 6, we identified 11 cell clusters, representing at least four cell types, including *POU5F1*^+^ pluripotent cells, *MESP1^+^* mesoderm, *ISL1*^+^ cardiac precursors and nascent *TNNT2*^+^ cardiomyocytes (Figure 2D). At day 11 and day 23, differentiated cell types were assigned and present in all genotypes (Figure 2E-H, S2A, B), based on cell type-specific marker genes (DeLaughter et al., 2016; Li et al., 2016). This included a diversity of *TBX5*^+^ cell types, comprising very few *PLVAP*^+^ endothelial cells or *TTR*^+^ endodermal cells, some *COL1A1*^+^ fibroblasts and, most abundantly, *TNNT2*^+^/*IRX4*^+^ ventricular cardiomyocytes (Figure 2E-G, S2A, B).

**Figure 2.**
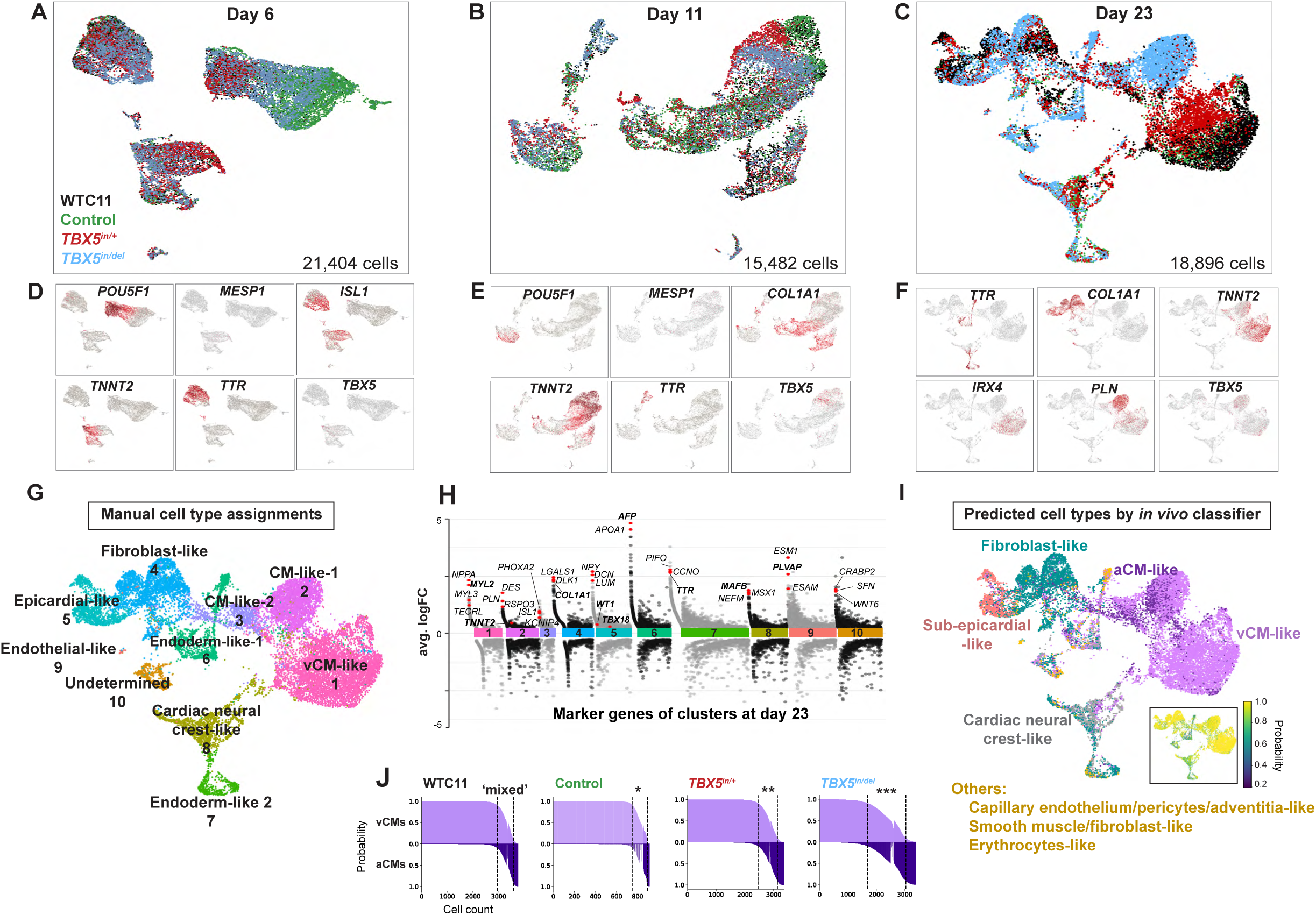
Human cardiomyocyte differentiation is sensitive to reduced TBX5 dosage. (A-C) Cells of each *TBX5* genotype were harvested at specific stages during directed differentiation to cardiomyocytes for single cell RNA-seq. UMAPs display cells of *TBX5* genotypes at day 6, day 11 or day 23. (D-F) Feature plots in UMAP space demonstrate expression of selected marker genes, which represent major cell types at each timepoint. (G) Cell type assignments of iPSC-derived cells at day 23 by manual annotation using marker genes are shown in UMAP space. (H) A Manhattan plot displays differentially expressed genes by cell type cluster at day 23. Examples of enriched genes by cluster are shown. Manual annotation was based on expression of bolded genes. (I) Predicted cell types of iPSC-derived cells at day 23 are classified using machine learning, based on human fetal cardiac cells *in vivo* (Asp et al., 2019). Inset of UMAP shows prediction probabilities for iPSC-derived cells at day 23 by the *in vivo* cell type classifier. (J) Waterfall plots for each *TBX5* genotype display prediction probabilities of iPSC-derived cells classified as ventricular cardiomyocytes (vCM), atrial cardiomyocytes (aCM) or mixed (<0.95 probability difference of vCM and aCM). * p<0.01, ** p<0.001, *** p<0.0001 by Fisher’s exact test.

We employed a machine learning algorithm (Pedregosa et al., 2011), to quantitatively evaluate the degree of similarity, if any, between iPSC-derived cells and cells from the developing human heart. A cell type classifier was trained on single cell gene expression from a human fetal four-chambered heart at 6.5-7 weeks gestation (Figure S2C) (Asp et al., 2019). This was used to predict cell type assignments for cells harvested at day 23 (Figure 2I, S2D, Table S1). Ventricular-like cardiomyocytes were the most commonly predicted cell type, constituting 52% of cells from all genotypes, with a high prediction probability average of 0.93, consistent with manual assignments by cell type-specific markers genes, such as *TNNT2* and *IRX4* (Figure 2G, I, S2B, D, E). Twenty-three percent of cells were assigned as fibroblast-like cells with 0.89 probability, 6% as epicardial-like cells with 0.89 probability, and 7% as cardiac cells of neural crest origin with 0.72 probability (Figure 2I, S2D, E). *AFP*^+^ or *TTR*^+^ cells, considered to be derived from endoderm or mesendoderm, were dispersed across several predicted cell assignments (Figure 2I, S2D). As expected, differentiation did not yield iPSC-derived cell types, including erythrocytes (1%) and immune cells (0.2%), which were sparsely represented with less than 0.5 prediction probability (Figure 2I, S2D). Taken together, the classifier’s predictions appeared to provide sufficient fidelity for assignments of iPSC-derived cells as *in vivo-*like cardiac cell types.

Although the cell type classifier was largely consistent with cell type assignments from manual annotations (Figure 2G, I, S2D), it predicted 9% of iPSC-derived cells from all genotypes as atrial-like cardiomyocytes with a 0.83 probability. Whereas the cell type classifier predicted a similar total number of high-probability (>0.7) cardiomyocytes for each *TBX5* genotype (Figure S2E), more atrial-like cardiomyocytes were predicted for *TBX5^inl+^* and *TBX5^in/del^* cells (p<0.0001 by Fisher’s exact test), than for WT (Figure S2E). The classifier also uncovered a population of iPSC-derived cardiomyocytes with ‘mixed’ identity, of both ventricular and atrial predictions. Interestingly, these ‘mixed’ cardiomyocyte predictions were more prevalent among *TBX5^in/+^* (p<0.001 by Fisher’s exact test) and *TBX5^in/del^* cells (p<0.0001), than wildtype or control (Figure 2J), supporting a notion that reduced *TBX5* dosage may perturb ventricular cardiomyocyte identity.

### TBX5 protects human ventricular cardiomyocyte fate

To assess if reduced TBX5 dosage disturbs paths of directed differentiation, we evaluated supervised URD trajectories from all *TBX5* genotypes and time points. URD predicts cellular chronology based on user-determined origins and ends (Farrell et al., 2018a). We defined *POU5F1*^+^ cells, which were predominantly from a single cluster at day 6 (Figure 2D), as the root and day 23 clusters as the tips in the pseudotime tree (Figure 3A, B). Cells at day 6 were found near the top of the tree, while cells at day 11 were distributed mid-way, followed by day 23 cells at the user-defined tips (Figure S3A-C). This demonstrated a logical ordering of cells along pseudotime by URD. Since *TBX5* transcripts were detected in all genotypes, including *TBX5^in/del^* cells, inferred lineage decisions for *TBX5*^+^ cell types in the absence of *TBX5* could be examined. We focused on inferred trajectories of *TBX5*^+^ cells to ventricular cardiomyocytes. *TBX5^in^*^/+^ cells followed a path similar to WT (Figure 3C, D, dashed lines), but to a transcriptionally distinct endpoint. In contrast, *TBX5^in^*^/*del*^ cells deviated from the WT differentiation path to ventricular cardiomyocytes (Figure 3E).

**Figure 3.**
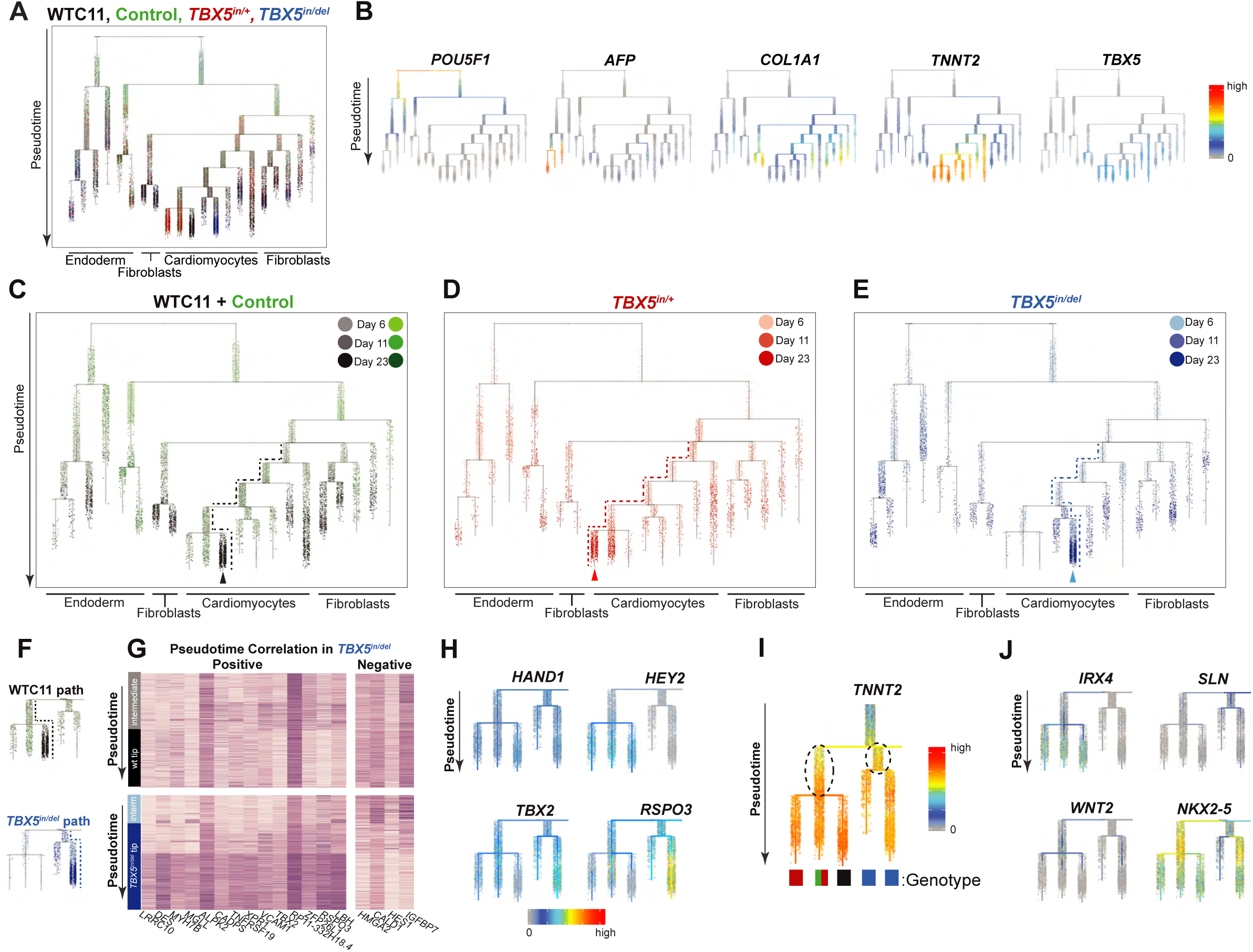
*TBX5* loss disturbs cell trajectories to ventricular cardiomyocyte fate. (A) Developmental trajectories were inferred by URD using a combined dataset of all *TBX5* genotypes and time points. A dendrogram shows cells at day 6, 11 and 23, during directed differentiation to cardiomyocytes. Pseudotime is displayed from root (top) to tips (bottom). Each *TBX5* genotype is color-coded from light to dark, to indicate the time point. (B) Expression of genes that define the major cell types (pluripotent cell, endoderm, fibroblast and cardiomyocyte) and *TBX5* are shown. (C-E) Cells are highlighted by *TBX5* genotype on the aggregate pseudotime dendrogram. Note the enrichment or depletion of cells from one genotype at certain branch points to tips (arrowheads). Dashed lines show a path to ventricular cardiomyocytes, by *TBX5* genotype. (F) Deduced paths to non-dividing cardiomyocytes of WT (black dashed line) or *TBX5^in/del^* (blue dashed line) are shown, from intermediates (labeled ‘interm’) to tips. (G) Heatmaps show expression for each gene that displays no correlation with pseudotime in the WT path (above), but a positive or negative correlation (|rho|≥0.4 and Z-score≥15 by difference in rho) in the *TBX5^in^*^/*del*^ path (below). (H) Feature plots show the WT or *TBX5^in^*^/*del*^ path for ventricular cardiomyocyte-enriched genes *HAND1* and *HEY2*, and atrioventricular canal-enriched genes *TBX2* and *RSPO3*. (I) Differential gene expression of inferred precursors for the cardiomyocyte branches (dashed ovals) show several genes that display altered gene expression (adj p-value<0.05 by Wilcoxon Rank Sum test) along the deduced WT or *TBX5^in^*^/*del*^ path. Colored blocks below the dendrogram represent the predominant *TBX5* genotypes in each tip. (J) The ventricular cardiomyocyte-enriched gene *IRX4* was absent along the *TBX5^in^*^/*del*^ path. *SLN* was qualitatively enriched in the *TBX5^in^*^/*del*^ path. Activation of *WNT2* and *NKX2-5* in the deduced *TBX5^in^*^/*del*^ path was delayed. Significance was determined by Wilcoxon Rank Sum test (adj p-value<0.05).

In order to explore gene expression changes that may have led to this deviation, we identified genes that change as a function of pseudotime in the WT or *TBX5^in^*^/*del*^ paths (Figure 3F). We deduced 22 genes (e.g. electrophysiology-related *NAV1* and *TECRL,* cardiomyopathy-associated *LAMA4,* and small peptide hormone *NPPA*), which were positively correlated with pseudotime in the WT/*TBX5^in/+^* branch (p-value<0.05 by two-sided *t* test), but aberrant in the *TBX5^in/del^* branch (Z-score≥15), suggesting that these genes were not activated properly in *TBX5^in/del^* cells (Figure S3D). Conversely, five genes were negatively correlated in the WT/*TBX5^in/+^* branch (p-value<0.05 by two-sided *t* test), but not in the *TBX5^in/del^* branch (Z-score≥15). Likewise, we identified 18 genes that positively (e.g. sarcomere *DES,* vascular adhesion *VCAM1,* and TF *LBH*) or negatively (e.g. TF *HES1* and actomyosin binding *CALD1*) correlated with pseudotime in *TBX5^in/del^* cells (p-value<0.05), but were altered in wildtype cells (Z-score≥15) (Figure 3G), signifying that these genes were inappropriately deployed in *TBX5^in/del^* cells.

A few ventricular markers (e.g. cardiac TFs *IRX4* and *HEY2*) were absent in *TBX5^in/del^* cells (Figure 3H, J), reminiscent of features from mouse (Bruneau et al., 2001). However, *TBX5^in/del^* cells still expressed other ventricular-enriched genes (e.g. cardiac TFs *HAND1* and *HAND2*) (Figure 3H), consistent with their electrophysiologic characteristics as beating ventricular cardiomyocytes (Figure 1J). In conjunction, *TBX5^in/del^* cells expressed markers of the atrioventricular canal (e.g. cardiac TF *TBX2* and Wnt agonist *RSPO3*) (Figure 3H), indicating that *TBX5* loss results in a disordered ventricular cardiomyocyte-like identity with ectopic gene expression.

We tested differential gene expression between intermediate branches, to identify genes that determine TBX5-dependent ventricular cardiomyocyte differentiation. We considered these branches as potential precursors proximal to *TBX5* genotype-specific tips. We compared these intermediate branches of cells that distinguish the cell trajectory route of WT and *TBX5^in^*^/+^ to *TBX5^in^*^/*del*^ (Figure 3I). These included secreted factors or cell surface receptors (*WNT2*, *FGFR1*; adj p-value<0.05) and cardiac TFs (*IRX4*, *HAND2*; adj p-value<0.05). Of note, expression of the CHD cardiac transcription factor *NKX2-5*, a transcriptional partner of TBX5 (Bruneau et al., 2001; Hiroi et al., 2001; Luna-Zurita et al., 2016), was differentially expressed between genotype-enriched intermediate branches of the URD tree (Figure 3J; adj p-value<1E-300 by Wilcoxon Rank Sum test). Consistent with a role of *Nkx2-5* for mouse ventricular cardiomyocyte specification *in vivo* (Lyons et al., 1995; Tanaka et al., 1999), onset of *NKX2-5* expression was delayed in *TBX5^in^*^/*del*^ cells (adj p-value<0.05 by Bonferroni-Holm multiple testing correction). In conjunction, a module of genes (chromatin regulator *PARP1,* ribosome *RPL37,* junctional protein encoding *KIAA1462* and Na^+^/K^+^ transport *ATP1A1*; adj p-value<0.05), were expressed concomitantly with *NKX2-5* (Figure S3E, F). This provides a potential molecular explanation for the observed delay in the onset of beating by *TBX5* loss (Figure 1D).

### Discrete transcriptional responses to reduced TBX5 dosage in cardiomyocytes

*TBX5* genotype-specific clusters emerged among cardiomyocytes at day 11 (Figure 2B), and *TBX5* genotype-specific segregation was more striking at day 23, particularly in *TNNT2*^+^ cells (Figure 2C). Therefore, we focused on *TNNT2*^+^ clusters at day 23 (Figure 4A, B). First, we used a low resolution for Louvain clustering to assess genes that are highly differential between *TBX5* genotype-driven *TNNT2^+^* clusters (Figure 4C). We detected 121 genes that were differentially expressed between WT and *TBX5^in^*^/+^-enriched clusters (Figure 4D, Table S2). Five hundred twenty genes showed differential expression between WT and *TBX5^in^*^/del^-enriched clusters (Figure 4E, Table S2). To identify stepwise *TBX5* dose-dependent genes, we evaluated genes that were differentially expressed between WT vs. *TBX5^in^*^/+^-enriched clusters and WT vs. *TBX5^in^*^/+^-enriched clusters. We found 85 genes common to both lists with a multitude of expression patterns (Figure 4F, Table S2). Many genes displayed changes in both expression level and percentage of expressing cells (e.g. small peptide hormone *NPPA,* Wnt agonist *RSPO3,* arrhythmia-linked *TECRL,* sarcomere *DES*) (Figure 4F). A few genes showed similar levels of gene expression, with changes to percentage of expressing cells (e.g. serine hydrolase *MGLL* or CHD TF *ANKRD1* in *TBX5^in^*^/+^-enriched clusters). Some genes, such as *NPPA,* were highly sensitive to TBX5 dosage, with reduced expression in *TBX5^in/+^* nearly comparable to that in *TBX5^in/de^*^l^. In contrast, *TECRL* was partly reduced in *TBX5^in/+^* cells and was further decreased in *TBX5^in/de^*^l^. Notably, some genes were altered in *TBX5^in/+^* cells but had elevated levels in *TBX5^in/de^*^l^ cells (e.g. *TBX5,* myosin light chain *MYL9,* cardiac TF *HOPX*, sarcomere *DES*). Specifically, *TBX5* expression likely reflected apparent upregulation of non-mutated exons in *TBX5^in/de^*^l^ cells, as seen in the mouse (Mori et al., 2006), although TBX5 protein expression was not detected (Figure 1B). We speculate that expression of other genes with potentially counterintuitive behavior, such as *DES*, may reflect a type of regulatory network compensation or overcompensation, or perhaps indicate a disordered cell type.

**Figure 4.**
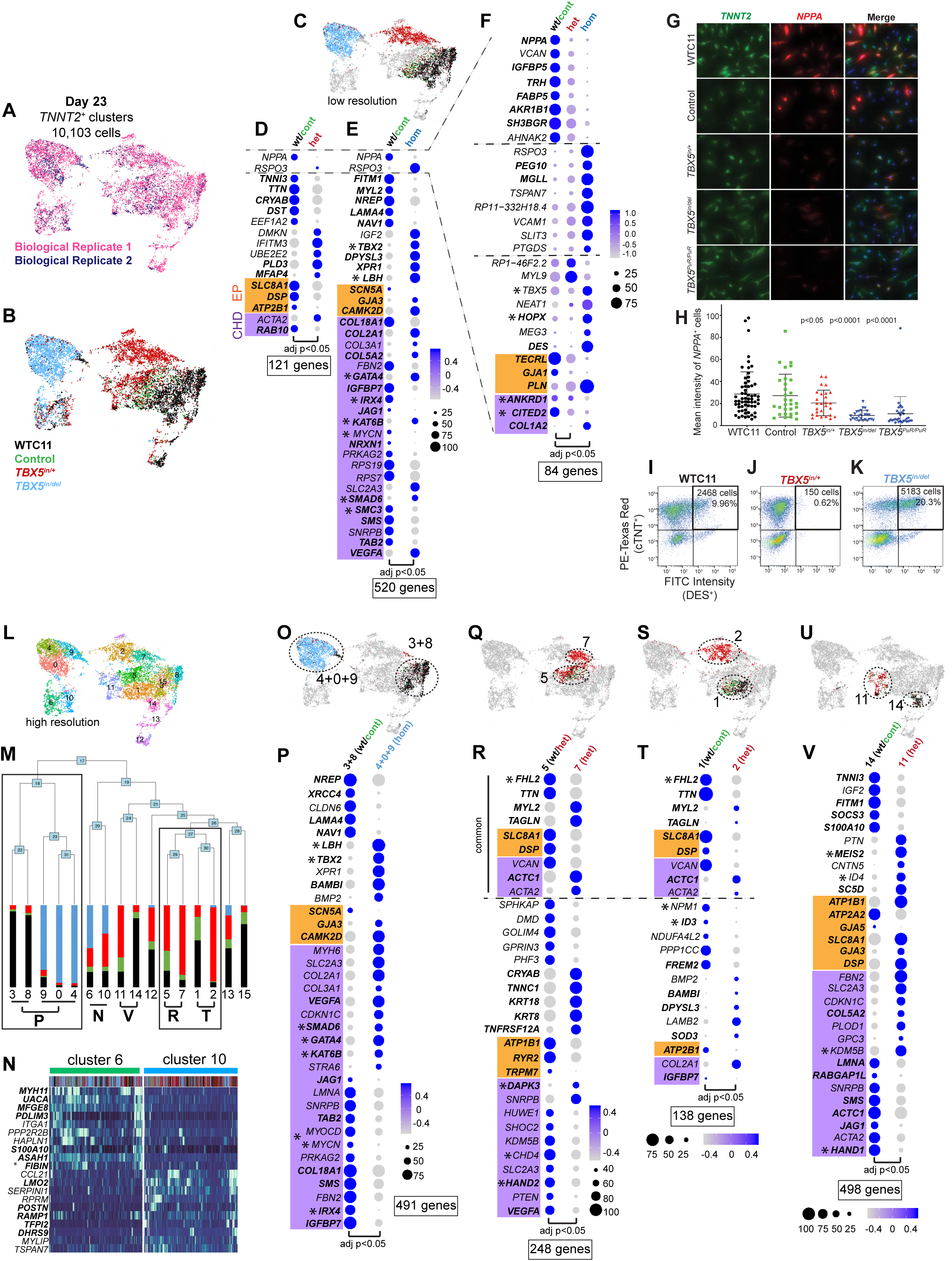
Subsets of cardiomyocytes respond discretely by quantitative transcriptional perturbations to reduced TBX5 dosage. (A-B) *TNNT2*^+^ clusters from day 23 were re-clustered in Seurat. UMAP shows cells colored by biological replicate (A) or *TBX5* genotype (B). (C) *TBX5* genotype-dominant clusters segregated at low resolution of Louvain clustering. (D) Clusters enriched for WT or *TBX5^in^*^/+^ were compared by differential gene expression. Top five upregulated or downregulated genes are displayed. EP (orange) or CHD (purple) genes are shown. Transcriptional regulators are denoted by asterisks, and predicted targets of TBX5, based on TBX5 occupancy (Ang et al., 2016) are bolded (Table S4). Dot size corresponds to the percentage of cells expressing the gene in a cluster, while the color intensity represents scaled expression values in a cluster. Significance was determined by Wilcoxon Rank Sum test (adj p-value<0.05). (E) Clusters enriched for WT or *TBX5^in^*^/d*el*^ were compared by differential gene expression. (F) Common genes that were differentially expressed between WT-vs. *TBX5^in^*^/+^-enriched clusters and WT vs. *TBX5^in/del^*-enriched clusters are shown. (G) Fluorescence *in situ* hybridization is visualized for *TNNT2* (green) or *NPPA* (red) in day 23 cardiomyocytes, from WTC11, control, *TBX5^in/+^*, *TBX5^in/del^* and *TBX5^PuR/PuR^* cells. Brightness and contrast of images have been adjusted to facilitate viewing of cells. (H) Graph displays mean intensity of *NPPA* signal of individual double-positive *TNNT2*^+^/*NPPA*^+^ cells by *TBX5* genotype. Significance of p-values were calculated by unpaired *t* test. (I-K) Pseudocolor plots of flow cytometry show cTNT^+^ or DES^+^ cells in (I) wildtype, (J) *TBX5^in/+^* or (K) *TBX5^in/del^* at day 23. The number of double-positive cTNT^+^/DES^+^ cells are significantly different between wildtype and *TBX5^in/+^* and between wildtype and *TBX5^in/del^* (p-value<1E-4 by Chi-Square test). (L) UMAP shows cells colored by cluster at higher resolution of Louvain clustering, indicating putative *TNNT2^+^* subsets. (M) A phylogenetic tree shows the relatedness of the ‘average’ cell in each cluster using PC space. The proportion of cells in each cluster are colored by *TBX5* genotype. Related clusters between different *TBX5* genotypes were selected for differential gene tests. (N) Heatmap shows hierarchically-sorted enriched genes in clusters 6 or 10, which consists of each *TBX5* genotype. (O) UMAP displays combined WT-enriched (black) or *TBX5^in/del^*-enriched (blue) clusters for comparison. (P) Dot plots show top five differentially expressed upgregulated or downregulated genes, along with EP or CHD genes, between aggregate WT-enriched and *TBX5^in/del^*-enriched clusters. (Q, S, U) UMAPs highlight clusters used for pair-wise comparisons for differential gene tests in corresponding dot plots below. (R, T, V) Dot plots of top differentially expressed genes between WT/*TBX5^in/+^*-enriched cluster 5 and *TBX5^in/+^*-enriched cluster 7 (R), between WT-enriched cluster 1 and *TBX5^in/+^*-enriched cluster 2, or (V) between WT-enriched cluster 14 and *TBX5^in/+^*-enriched cluster 11. A few differentially expressed genes are common between comparisons in (R) and (T). Total number of differentially expressed genes for each comparison is listed (Table S2).

We used orthogonal assays at single cell resolution to validate examples of TBX5-dependent genes. *TBX5* dosage-dependent downregulation of *NPPA* was evident in cardiomyocytes by RNAscope (*TBX5^in/+^*, p<0.05; *TBX5^in/de^*^l^ or *TBX5^PuR/PuR^*, p<1E-4 by Student’s t-test) (Figure 4G, H), consistent with the TBX5-dependent rheostatic regulation of *Nppa* in mouse (Bruneau et al., 2001; Mori et al., 2006). By flow cytometry, DES protein was reduced in *TBX5^in/+^* (p-value<1E-4 by Chi-Square test) and upregulated in *TBX5^in^*^/*del*^ (p-value<1E-4 by Chi-Square test) cardiomyocytes, compared to wildtype (Figure 4I-K), corroborating this pattern of *TBX5* dose-dependent expression.

To assess the heterogeneity among cardiomyocytes, we used a higher resolution for Louvain clustering and constructed a phylogenetic cluster tree relating 16 different *TNNT2*^+^ cell clusters (Figure 4L, M). We considered these clusters as putative functional subpopulations of ventricular cardiomyocytes, since they could not be classified based on a conventional anatomy-based categorization. We found two clusters (clusters 6 and 10) that included a similar proportion of cells from each *TBX5* genotype, implying that these putative cardiomyocyte subpopulations may be insensitive to reduced TBX5 dosage (Figure 4M, N). We then searched for differentially expressed genes by pairwise comparisons of related subpopulations between *TBX5* genotypes (Figure 4O-U). For example, cluster 5 contains WT and *TBX5^in^*^/+^ cells (Figure 4Q), suggesting that these *TBX5* heterozygous cells are indistinguishable from a subpopulation of WT. In contrast, cluster 7 is largely composed of *TBX5^in^*^/+^, suggesting that these TBX5 heterozygous cells are distinct. In addition to stepwise TBX5 dose-dependent genes (Figure 4F) that were often altered in many cluster-to-cluster comparisons, we detected additional common changes in gene expression amongst pairwise cluster comparisons of WT vs. *TBX5^in^*^/+^ clusters. These included the cardiac TF *FHL2*, the cardiomyopathy-linked sarcomere gene *TTN*, and the ventricular-enriched sarcomere gene *MYL2* (Figure 4Q-V; adj p-value<0.05 by Wilcoxon Rank Sum test). We also discerned many differences in gene expression based on cluster-specific comparisons (adj p-value<0.05 by Wilcoxon Rank Sum test), implying varied transcriptional responses among subpopulations of cardiomyocytes to *TBX5* haploinsufficiency (Figure 4Q-V) or *TBX5* loss (Figure 4O, P).

These differentially expressed gene sets at day 23 were enriched for electrophysiology (EP) genes (FDR<0.05, Figure 4, Table S2-4), which are implicated in membrane depolarization (*SCN5A*), calcium handling (*RYR2*, *ATP2A2*, and *PLN*) and arrhythmias (*TECRL*) (Figure 4). These genes provide a molecular explanation for the EP defects observed upon *TBX5* mutation. Several altered transcripts were encoded by candidate genes implicated in CHD (e.g. TFs *CITED2, MYOCD,* and *ANKRD1*) (Figure 4, Table S2-4) (Homsy et al., 2015; Jin et al., 2017; Lalani and Belmont, 2014; McCulley and Black, 2012; Prendiville et al., 2014; Priest et al., 2016; Sifrim et al., 2016; Zaidi et al., 2013). In addition, some TBX5-dependent genes that were previously associated with CHD or arrhythmias by genome-wide association studies (GWAS) were identified (Cordell et al., 2013a; 2013b; Ellinor et al., 2012; Hoed et al., 2013; Hu et al., 2013; Pfeufer et al., 2010a; Smith et al., 2011). We uncovered *IGFBP7, MYH7B* and *SMCHD1* for CHD and 45 reported genes for arrhythmias (for example, *PLN, HCN4, SCN5A, GJA1*, *PITX2* and *TECRL*; FDR<0.05) among TBX5-sensitive genes (Table S2-4).

We assessed if TBX5 dose-sensitive genes were largely direct or indirect targets of TBX5, by examining TBX5 occupancy in human iPSC-derived CMs from a published dataset (Ang et al., 2016). We found correlations of TBX5 occupancy near TBX5 dosage-vulnerable gene sets at day 23 (Figure 4, bolded genes; Figure S4A-F, Table S4). For example, 61 of 85 genes that showed stepwise dose-dependence were near TBX5 binding sites (Figure 4F, Figure S4A), suggesting that these genes were predicted targets of TBX5. TBX5 cooperates with GATA4 for cardiac gene regulation (Ang et al., 2016; Garg et al., 2003; Luna-Zurita et al., 2016). We also observed a high association of GATA4 occupancy with TBX5 (Ang et al., 2016) near TBX5-dependent genes (Figure S4A-E, Table S4, 5), indicating that GATA4 may have a role in modulating TBX5 dosage-sensitive genes.

Since modifiers in different genetic backgrounds can modulate phenotypic effects, we assessed alternatively targeted iPSC lines of *TBX5* mutants in an independent genetic background (PGP1, from a Caucasian male (Lee et al., 2009), compared to WTC11 from a Japanese male (Miyaoka et al., 2014), Figure S5A, B). We independently evaluated comparisons between genotype-enriched subtype clusters in day 23 *TNNT2*^+^ cells from PGP1-derived cell lines (Figure S5C-G). Comparisons of lists of TBX5-dependent genes in day 23 *TNNT2*^+^ cells showed overlap between WTC11 vs. *TBX5^in/+^* and PGP1 vs. *TBX5^in/+^* (p<5.219e-81 by hypergeometric test), or WTC11 vs. *TBX5^in/del^* and PGP1 vs. *TBX5^del/del^* (p<1.438e-172).

We also integrated day 23 *TNNT2*^+^ cells from each genetic background into one combined dataset for analysis. Cells were largely indistinguishable in UMAP space regardless of experimental replicate or genetic background (Figure S5H). Importantly, we again observed segregation by *TBX5* genotypes (Figure S5I). By comparing genotype-enriched subtype clusters (Figure S5J, K), we detected 148 genes between WTC11/Control/PGP1 and *TBX5* heterozygous cells, and 457 genes between WTC11/Control/PGP1 and *TBX5* homozygous cells (Figure S5L, M, Table S2). These results demonstrated robust TBX5 dosage-dependent gene expression alterations in cardiomyocytes from independent experiments, genetic backgrounds, and gene targeting strategies. Any differences in gene expression between biological replicates and genetic backgrounds likely reflected a combination of technical variability, biological stochasticity or genetic modifiers that, as in patients with *TBX5* mutations (Basson et al., 1994), may explain variable expressivity of disease for a given mutation.

### TBX5 dosage maintains cardiac gene network stability

CHD-associated and arrhythmia-related genes were enriched among TBX5-dependent genes in complex patterns of expression. We sought to independently, and without bias, assess the importance of TBX5 in a global cardiac gene regulatory network (GRN) beyond changes to gene expression. To evaluate the role of TBX5 dosage for regulating GRNs, we used bigSCale2 (Iacono et al., 2019) to independently infer putative GRNs without *a priori* knowledge (e.g. protein-protein interactions, known genetic associations, cardiac-enriched genes) from single cell expression data of *TNNT2*^+^ cells. By applying the concept of “pagerank”, first devised to rank the importance by popularity of websites via numerical weighting (Brin and Page, 1998), we predicted quantitatively the biological importance (i.e. centrality) of genes in a GRN, even if a node’s gene expression was unchanged by *TBX5* dosage.

By comparing inferred networks of WTC11 and Control to *TBX5^in/+^* or *TBX5^in/del^* within a given time point, we uncovered several candidate nodes that displayed loss of pagerank centrality from reduced TBX5 dosage (Figure 5A-C, S6A-D Table S6). These included the calcium-handling gene *RYR2*, and twenty CHD genes (for example, TFs *GATA6*, *HAND2*, and *SMAD2,* p<2.2e-5 by hypergeometric test,) (Figure 5C), consistent with our analysis from differential gene expression. For example, at day 11, pagerank centrality of the CHD TF *SMAD2* was absent in *TBX5^in/+^* cells (Figure 5A-C, top 5% of all changes), indicating a possible impairment of *SMAD2* function from *TBX5* haploinsufficiency. Centrality of the cardiac development-related TF *MEF2C*, which is necessary for mouse heart development (Lin et al., 1997), was substantially reduced by heterozygosity or loss of *TBX5* at day 11 (Figure 5A-C, top 5% cutoff). Quantitative alterations to GRNs showed that TBX5 dosage may be critical for maintaining cardiac network stability, and potentially unveiled putative genetic interactions disrupted in TBX5-dependent CHDs.

**Figure 5.**
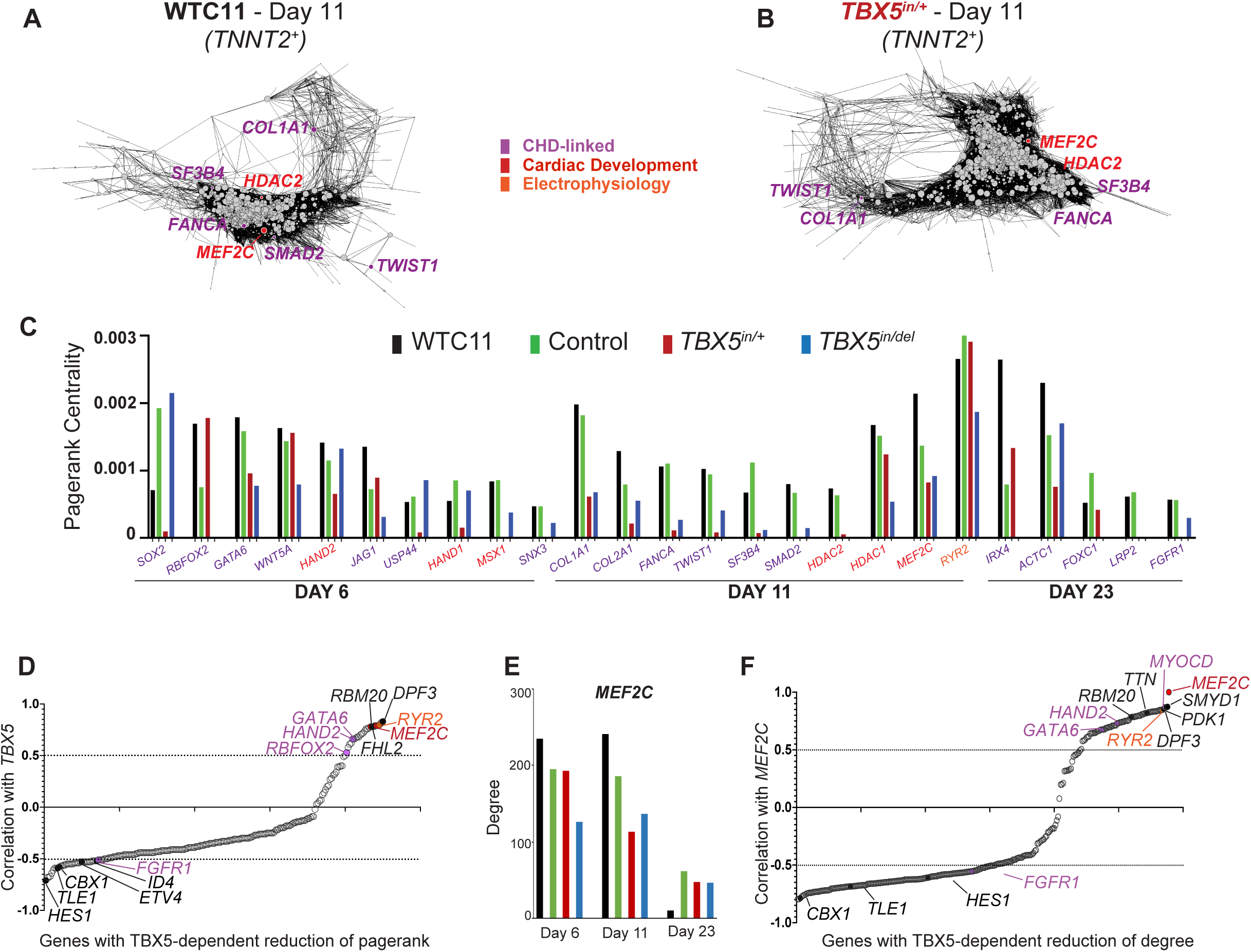
TBX5 dosage preserves cardiomyocyte network stability. (A, B) Gene regulatory networks (GRNs) of *TNNT2^+^* cells for each *TBX5* genotype at day 6, 11, or 23 were inferred. GRNs at day 11 for WTC11 (A) or *TBX5^in/+^* (B) are shown. Nodes of CHD (purple), heart development (red) or electrophysiology (orange) genes are shown. The size of each node represents the quantitative importance of the gene, based on pagerank centrality. Note the absence of *SMAD2* and the reduced centrality of *MEF2C* (smaller circle) in the *TBX5^in^*^/+^ network, compared to WTC11. (C) Pagerank centrality for significantly altered (top 5% cutoff) nodes of CHD, heart development or EP genes at specific time points are shown. Twenty CHD genes display a reduction in pagerank (top 5% cutoff, when compared to wildtype and control) in at least one *TBX5* mutant genotype at any stage. This indicates enrichment of CHD genes in TBX5 dosage-sensitive networks (p<2.2e-5 by hypergeometric test). (D) TBX5-dependent genes with a reduction of pagerank are correlated (correlation >0.5), anti-correlated (correlation <-0.5), or indeterminate (0.5<correlation<-0.5) with *TBX5* expression in *TNNT2^+^* cells. (E) Degree centrality for *MEF2C* is reduced in *TBX5^in/del^* at day 6 and reduced in *TBX5^in/+^* and *TBX5^in/del^* at day 11 (top 5% cutoff, when compared to wildtype and control), but not at day 23. (F) Correlations with *MEF2C* and TBX5-dependent genes with a reduction of degree centrality in *TNNT2^+^* cells are plotted. Additional data can be found in Table S6.

To further investigate the predicted relationship between *TBX5* and *MEF2C* within a human TBX5 dosage-sensitive, CHD-associated GRN, we used a complementary approach using bigSCale2, to identify gene-gene correlations with *TBX5* expression in individual *TNNT2*^+^ cells across timepoints and *TBX5* genotypes (Figure 5D). Genes highly co-expressed with *TBX5* (Pearson coefficient >0.5) regardless of *TBX5* genotype suggested potential positive regulation or possible cell autonomous effects by *TBX5* dosage (for example, calcium-handling *PLN* and *RYR2*, and sarcomere *TTN*), while those with high anti-correlation (Pearson coefficient <-0.5) suggested potential negative regulation or possibly non-cell autonomous effects (for example, TFs *HES1, TLE1, CBX1, ETV4, ID4,* and cell surface receptor *FGFR1*). *MEF2C* expression was among the highest correlated with *TBX5* expression and demonstrated the greatest TBX5-dependent decrease of pagerank at day 11 (Figure 5D, Table S6), further suggesting *MEF2C* as a putative candidate for mediating TBX5 dose-sensitive regulatory effects.

*MEF2C* gene expression itself was unchanged by reduced TBX5 dosage. Yet, *MEF2C* also displayed the greatest TBX5-dependent decrease in degree centrality, which reflects a node’s connections in a network and contributes to pagerank, at day 11 (Figure 5E, Table S6). This indicated potential alterations to *MEF2C* functional connectivity within the TBX5-dependent GRN. We found that multiple genes, which correlated with *MEF2C,* displayed diminished levels of degree by reduced TBX5 dosage (e.g. transcriptional regulators *SMYD1* and *MYOCD,* sarcomere *TTN,* calcium-handling *RYR2,* and kinase *PDK1*; top 5% cutoff) (Figure 5F, Table S6). Some genes (*SMYD1* and *MYOCD*) are direct MEF2C targets in mice *in vivo* (Creemers et al., 2006; Phan et al., 2005). This suggested to us that these candidate genes with reduced degree may mediate putative *MEF2C* functional connectivity for TBX5 dosage-sensitive GRNs.

### *Tbx5* and *Mef2c* cooperate for ventricular septation *in vivo*

Several potential genetic interactions were predicted by reduced pagerank from TBX5 dose-dependent human GRNs. A predicted genetic interaction between *Tbx5* and *Gata6* is known from mouse studies (Maitra et al., 2009). However, heterozygous loss of *Tbx5* can lead to highly penetrant perinatal lethality based on mouse genetic background strains (Bruneau et al., 2001; Mori et al., 2006), making it difficult to evaluate genetic interactions based on postnatal lethality. Therefore, we further characterized a multifunctional allele of *Tbx5* (*Tbx5^CreERT2IRES2xFLAG^*, abbreviated *Tbx5^CreERT2^*) (Devine et al., 2014a), which appeared to be a hypomorphic *Tbx5* allele, as a potential genetic tool for probing highly-sensitive *in vivo* genetic interactions with *Tbx5* (Figure 6). Mice heterozygous for *Tbx5^CreERT2IRES2xFLAG^* (*Tbx5^CreERT2/+^*) survived to adulthood, and Mendelian ratios were recovered at weaning, as well as during embryonic development (Figure 6A). However, embryos homozygous for *Tbx5^CreERT2IRES2xFLAG^* (*Tbx5^CreERT2/CreERT2^*) could only be recovered until embryonic day 16.5 (E16.5), indicating that the *Tbx5^CreERT2IRES2xFLAG^* allele is likely hypomorphic (Figure 6B). Histological analysis of embryonic *Tbx5^CreERT2/CreERT2^* hearts at E16.5 showed atrioventricular canal (AVC) defects, which include atrial septal defects (ASDs), VSDs and an atrioventricular valve (AVV), which were not present in wildtype or *Tbx5^CreERT2/+^* mice, implicating CHDs as the cause of late embryonic lethality (Figure 6C - F).

**Figure 6.**
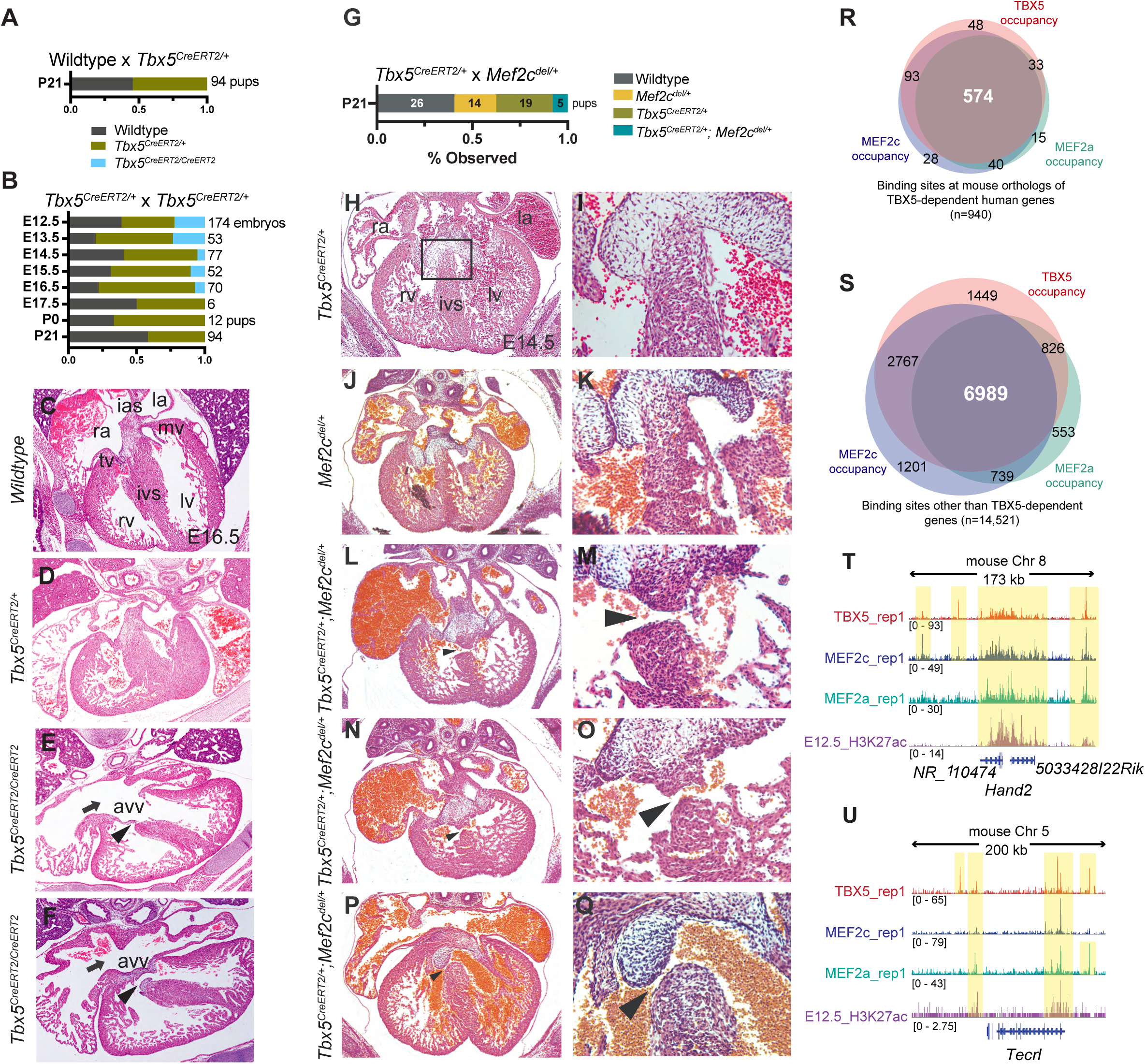
*Tbx5* and *Mef2c* cooperate in heart development. (A) Pups at postnatal day 21 (P21) from matings of wildtype X *Tbx5^CreERT2/+^* were genotyped, and expected Mendelian ratios were observed. (B) Expected Mendelian ratios from *Tbx5^CreERT2/+^* X *Tbx5^CreERT2/+^* were not observed for *Tbx5^CreERT2/CreERT2^* embryos after embryonic day 13.5 (E13.5), and none were recovered beyond E16.5. (C-F) Transverse sections of hearts at E16.5 from each *Tbx5* genotype are shown. In *Tbx5^CreERT2/CreERT2^* embryos, note atrioventricular canal defects, consisting of atrioventricular valves (AVV), ventricular septal defects (arrowhead), and atrial septal defects (arrow). LV, left ventricle; RV, right ventricle; IVS, interventricular septum; LA, left atrium; RA, right atrium; IAS, interatrial septum. (G) Pups at P21 from matings of *Tbx5^CreERT2/+^* X *Mef2c^del/+^* were genotyped. Expected Mendelian ratios were not observed for *Tbx5^CreERT2/+^;Mef2c^del/+^* at P21. (H, J, L, N, P) Transverse sections of hearts at embryonic day 14.5 (E14.5) from each genotype are shown. (I, K, M, O, Q) Magnified views of the interventricular septum are shown. In *Tbx5^CreERT2/+^;Mef2c^del/+^* embryos, VSDs are observed, including muscular VSDs (arrowheads in M, O), a subaortic membranous VSD (Q, arrowhead) and dilated blood-filled atria (L, N, P). (R, S) Venn diagrams display the overlap of TBX5, MEF2a or MEF2c occupancy near mouse orthologs of human TBX5-dependent (R) or - independent genes (S). (T, U) Browser tracks for ChIP-seq data from E12.5 hearts for TBX5, MEF2c, MEF2a and H3K27ac near TBX5-dependent genes, *Hand2* (T) or *Tecrl* (U). Yellow bands of shading indicate co-occupancy. Table S7 displays odds ratios (FDR<0.05) of TBX5, MEF2a or MEF2c occupancy near human TBX5-dependent genes, while Table S8 displays odds ratio (FDR<0.05) of co-occupancy of TBX5, MEF2a and MEF2c near mouse orthologs of TBX5-dependent human genes.

*MEF2C* was a quantitatively important node in human TBX5-dependent GRNs. Accordingly, we evaluated a predicted genetic interaction between *TBX5* and *MEF2C* in an *in vivo* mammalian context. Using the hypomorphic allele of *Tbx5* and a null allele of *Mef2c (Mef2c^del^)* (Lin et al., 1997), we noted that *Tbx5^CreERT2/+^;Mef2c^del/+^* mice were underrepresented at weaning (Figure 6G). By histology, we detected a highly penetrant morphologic phenotype of ventricular septal defects (n=4 of 4), consisting of muscular (n=3 of 4) or membranous (n=1 of 4) VSDs, in compound heterozygous embryos at E14.5. VSDs were not observed in *Tbx5^CreERT2/+^* or *Mef2c^del/+^* littermate embryos (Figure 6H-Q). Muscular VSDs are rarely observed in mouse models of CHD, making this observation particularly compelling. These findings demonstrate a highly-sensitive genetic interaction between *Tbx5* and *Mef2c* in mouse *in vivo*, consistent with predictions from a human TBX5 dose-sensitive GRN.

We speculated that MEF2C may play a direct role to co-regulate TBX5-dependent gene expression during heart development. Using mice targeted with a FLAG-biotin (fl-bio) tag at specific TF loci, chromatin occupancy (Akerberg et al., 2019) of TBX5, MEF2C, and MEF2A (also predicted to be part of the TBX5-dependent GRN, Table S6) was highly correlated near mouse orthologs of TBX5-sensitive human genes (for example, *HAND2*, *FHL2*, *TECRL*, *NPPA*/*NPPB*; Figure 6R-U; Tables S7-8, FDR<0.05 for multiple comparisons). Thus, direct co-regulation of target genes by TBX5, MEF2C, and MEF2A, in addition to previously known co-occupancy with NKX2-5 and GATA4 (Ang et al., 2016; Luna-Zurita et al., 2016), may be a potential TBX5 dosage-dependent mechanism for *TBX5* haploinsufficiency.

## Discussion

Our studies with a human cellular model of *TBX5* haploinsufficiency has defined consequences of reduced TBX5 dosage during cardiomyocyte differentiation at single cell resolution, indicating a dose-sensitive requirement of *TBX5* for human ventricular cardiomyocyte differentiation and function. Of potential relevance to a range of anatomical and functional manifestations of *TBX5* haploinsufficiency, we uncovered discrete responses to reduced TBX5 dosage in susceptible ventricular cardiomyocyte subsets. The quantitative specificity of TBX5-dependent cell types underscores cellular complexity in response to reduced transcription factor dosage. Many of the cellular phenotypes of this human disease model are cardiomyocyte-specific, intrinsic and likely cell autonomous. Dysregulated gene expression of EP or CHD genes provide potential molecular explanations for these cellular phenotypes, which are relevant to HOS, and more broadly to CHDs, in humans.

We found that *TBX5* dosage was necessary for preserving ventricular cardiomyocyte identity. We leveraged machine learning to predict assignments of iPSC-derived cells from a classification of human fetal heart cell types (Asp et al., 2019), lending support to the notion that our human disease modeling may serve as a reasonable proxy to study human cardiogenesis using molecular genetics. We predicted a mixed ventricular-atrial cardiomyocyte identity *in vitro*, which was exacerbated by *TBX5* loss. In addition, developmental trajectory inferences highlighted how a path to ventricular cardiomyocyte fate was vulnerable to reducing TBX5 dosage.

Susceptibility to TBX5 dosage-dependent gene expression in specific regions of the developing heart was apparent from studies modeling *TBX5* haploinsufficiency in the mouse (Bruneau et al., 2001; Mori et al., 2006). The implication would be that discrete populations in the developing human heart would respond specifically to reduced TBX5 dosage. In support of this notion, with single cell resolution of gene expression in human iPSC-derived cardiomyocytes, we detected discrete changes to reduced *TBX5* dosage in apparent subpopulations of human ventricular cardiomyocytes. The richness of detail achieved here eclipses current knowledge of *TBX5* haploinsufficiency from mouse models.

Many TBX5-sensitive genes that we discovered are related to heart function. CHDs are largely viewed as three-dimensional structural defects, but they are often accompanied by cardiac dysfunction, even after surgical correction. Arrhythmias and diastolic dysfunction are observed in patients with HOS (Basson et al., 1994; McDermott et al., 2008; Mori and Bruneau, 2004; Zhu et al., 2008), and in many other types of CHDs not related to *TBX5* (Panesar and Burch, 2017). Furthermore, TBX5 is strongly associated with EP defects based on genome-wide association studies (Ellinor et al., 2012; Pfeufer et al., 2010b; Smith et al., 2011). TBX5 dosage has been shown to be necessary for preserving diastolic function in mice, by modulating SERCA2a-dependent calcium transients (Zhu et al., 2008), and regulating calcium cycling in atrial myocytes in the context of atrial fibrillation (Dai et al., 2019; Laforest et al., 2019; Nadadur et al., 2016). In our iPS cell model, cardiomyocytes showed TBX5 dose-sensitive slowing of decay of calcium transients and disarray of sarcomeres, likely reflecting impaired ventricular cardiomyocyte relaxation. In these cells, dysregulation of several genes responsible for calcium cycling, including the SERCA2-encoding gene *ATP2A2*, NCX1-encoding gene *SLC8A1, RYR2*, and *PLN*, provide a potential molecular explanation for ventricular cardiomyocyte impairment and diastolic dysfunction in HOS (Eisner et al., 2020). Notably, the arrhythmia-associated gene *TECRL* displayed stepwise dosage-dependent sensitivity to reduced *TBX5* and is a predicted TBX5 target. Loss of *TECRL* in human iPSCs leads to prolonged calcium transients (Devalla et al., 2016), comparable to reducing TBX5 dosage. An understanding of TBX5 function in calcium homeostasis may uncover new mechanisms for human arrhythmogenesis and potentially for ventricular cardiomyocyte relaxation.

Many CHD genes were found to be altered due to reduced TBX5 dosage. This implies an interconnected network of CHD genes, potentially modulating each other’s functional targets. To gain an unbiased view into potential TBX5-sensitive networks, we inferred TBX5-dependent GRNs from individual human *TNNT2+* cells during a differentiation time course, from nascent to beating cardiomyocytes. We measured quantitative metrics for nodes of centrality, such as pagerank and degree (Iacono et al., 2019), and evaluated changes to each node by *TBX5* genotype and time point. Importantly, these quantitative measures of centrality are not defined by any *a priori* knowledge of cardiac biology, and stem solely from the single cell RNA-seq data. Quantitative analysis of human TBX5 dose-sensitive GRNs predicted vulnerable nodes enriched for CHD or cardiac development genes, suggesting a vital role for TBX5 dosage to maintain cardiac network stability. The sensitivity of a GRN to transcription factor dosage has been observed in *Drosophila* embryo patterning (Stathopoulos and Levine, 2002), for example, but has not been linked to human disease to date.

From TBX5-sensitive GRNs, we discovered several important nodes linking many CHD genes. For example, reduced centrality of *MEF2C* in the TBX5-dependent GRN predicted an important and sensitive genetic link between these cardiac transcription factors. Consistent with this notion, double-knockdown of *tbx5* and *mef2c* in zebrafish lead to severe defects in the looping heart tube (Ghosh et al., 2009). We observed a strikingly sensitive genetic interaction of *Tbx5* and *Mef2c* in mice using a hypomorphic allele of *Tbx5* (Devine et al., 2014b) and a null allele of *Mef2c* (Lin et al., 1997). This genetic interaction unveiled a finely tuned role later in mammalian heart development, beyond heart looping and chamber formation, for the process of ventricular septation. Of note, the *Tbx5* and *Mef2c* genetic interaction in mouse yielded muscular VSDs, a very specific type of CHD that is rarely observed in mouse models but common in humans. We anticipate that these and other genetic interactions will allow the discovery of molecular pathways and cellular processes that underlie specific CHDs.

While many TBX5-dependent genes were consistent across two ethnically diverse genetic backgrounds, there were some apparent differences. This is consistent with a notion that modifiers in genetic backgrounds can contribute to varying degrees of phenotypic expressivity for CHDs. Furthermore, variability in CHDs with monogenic inherited or *de novo* mutations could be explained by additional mutations or copy number variations of genes that form part of these functional regulatory networks, as illuminated by our findings, and as evidenced by oligogenic inheritance of CHD-causing variants (Gifford et al., 2019). Our results point to a genomic framework that will guide genetic insights into the underpinnings of CHD. The biophysical rules relating to transcription factor binding and dosage sensitivity are only now becoming understood. Our results in a human cellular model of *TBX5* haploinsufficiency may potentially bring immediate pertinence of human disease to this biological context.

## Contributions

I.S.K. and B.G.B conceived and designed the project. B.I.G., L.W., L.B., T.S. and I.S.K. performed gene targeting and isolation of mutant iPSCs. B.I.G., K.S.R, P.G., T.S., and I.S.K. performed in vitro differentiation and harvested samples. P.G. performed the Western analysis. M.H.L. performed electrophysiology analyses. R.T. performed statistical analyses for electrophysiology. K.S.R. performed immunostaining and scoring of cardiomyocytes. G.A.A. performed RNAscope and flow cytometry. K.S.R., A.P.B. and I.S.K. performed Seurat analysis. K.S.R., A.P.B., H.Z.G., and I.S.K. performed pseudotime analyses. A.P.B. employed machine learning for the cell type classifier. H.Z.G. implemented the cell browser. G.I. performed gene regulatory network analyses. W.P.D. and I.S.K. performed phenotype analyses of mutant mice. B.N.A., F.G., K.L., and W.T.P. performed ChIP-seq experiments and peak calling. S.K.H. and R.T. performed association analyses of co-occupancy, gene expression and disease candidates. J.M.S., W.T.P., C.E.S., J.G.S., and H.H. provided advising. I.S.K. and B.G.B. wrote the manuscript, with comments and contributions from all authors.

## Acknowledgements

We thank Dario Miguel-Perez and Sarah Wood for mouse genotyping and colony maintenance, Jeff Farrell for input on URD, David Joy and Todd McDevitt for sharing in-house imaging software, Brian Black for mouse lines, Kathryn Claiborn for editorial assistance, and members of the Bruneau lab for discussions and comments. We also thank the Gladstone Bioinformatics, Genomics, Histology and Microscopy, and Stem Cell Cores, the UCSF Laboratory for Cell Analysis, the UCSF Center for Advanced Technology, the Salk Institutes Center of Excellence for Stem Cell Genomics, Matthew Speir and Maximilian Haeussler at cells.ucsc.edu, and the UCSC Stem Cell Data Center Hub for their invaluable assistance. This work was supported by grants from the National Institutes of Health (NHLBI Bench to Bassinet Program UM1HL098179 to B.G.B. and UM1HL098166 to J.G.S., C.E.S. and W.T.P.; R01HL114948 to B.G.B., USCF CVRI 2T32HL007731-27 to S.K.H), the California Institute for Regenerative Medicine (RB4-05901 to B.G.B), the Office of the Assistant Secretary of Defense for Health Affairs through the Peer Review Medical Research Program under Award No. W81XWH-17-1-0191 (B.G.B), the Foundation for Anesthesia Education and Research (Mentored Research Training Grant to I.S.K.), Society for Pediatric Anesthesia (Young Investigator Award to I.S.K.), Hellman Family Fund (I.S.K.), UCSF REAC Grant (I.S.K.) and UCSF Department of Anesthesia and Perioperative Care (New Investigator Award to I.S.K.). H.H. is a Miguel Servet (CP14/00229) researcher supported by the Spanish Institute of Health Carlos III (ISCIII) and Ministerio de Ciencia, Innovación y Universidades (SAF2017-89109-P; AEI/FEDER, UE). This work was also supported by an NIH/NCRR grant (C06 RR018928) to the J. David Gladstone Institutes, and the Younger Family Fund (B.G.B.). Opinions, interpretations, conclusions and recommendations are those of the author and are not necessarily endorsed by the Department of Defense. In conducting research using animals, the investigator(s) adheres to the laws of the United States and regulations of the Department of Agriculture. In the conduct of research utilizing recombinant DNA, the investigator adhered to NIH Guidelines for research involving recombinant DNA molecules.

## METHODS

### CONTACT FOR REAGENT AND RESOURCE SHARING

All unique/stable reagents generated in this study are available from the Lead Contact, Benoit Bruneau with a completed Materials Transfer Agreement.

### EXPERIMENTAL MODEL AND SUBJECT DETAILS

#### Gene targeting and genotyping of human iPS cells mutant for *TBX5*

sgRNAs for *TBX5* exon 3 (sgRNA1, TCCTTCTTGCAGGGCATGGA) or exon 7 (sgRNA2, CCTTTGCCAAAGGATTTCG), which encode the T-box domain, were selected using crispr.genome-engineering.org, and cloned by annealing pairs of oligos into a plasmid containing humanized *S. pyogenes* Cas9, as described in (Cong et al., 2013) (px330-U6-Chimeric_BB-CBh-hSpCas9 was a gift from Feng Zhang, Addgene #42230).

For WTC11-derivatives *TBX5^+/+^* (control), *TBX5^in^*^/+^ or *TBX5^in^*^/*del*^, the induced pluripotent stem (iPS) cell line WTC11 (gift from Bruce Conklin, available at NIGMS Human Genetic Cell Repository/Coriell #GM25236) (Miyaoka et al., 2014) was electroporated (Lonza #VPH-5012) with a cloned nuclease construct containing a guide RNA (sgRNA1) targeting exon 3 of *TBX5*, as described in (Mandegar et al., 2016; Miyaoka et al., 2014). Cells were plated on human ESC-grade Matrigel (Corning #354277) and cultured in mTeSR-1 (StemCell Technologies Cat #05850) with 10µM ROCK inhibitor (StemCell Technologies, Y-27632). For screening of *TBX5* exon 3 non-homologous end-joining (NHEJ) mutations, genomic DNA flanking the targeted sequence was amplified by PCR (For1: ATGGCATCAGGCGTGTCCTATAA and Rev1: CCCACTTCGTGGAATTTTAGCCA), amplicons underwent digestion by NlaIII, and then were evaluated for loss of NlaIII by gel electrophoresis (wildtype band 800bp, mutant band 880bp). Clones with no change, a heterozygous or homozygous loss of NlaIII were sequenced (For1: ATGGCATCAGGCGTGTCCTATAA, Rev1: TTCCGGGCTTGAACTTCTGG, Seq1: ATAGCCTTGTGCTGATGGCA).

For generation of *TBX5^PuR/PuR^*, a puromycin resistance gene cassette (Frt-PGK-EM7-PuroR-bpA-Frt) containing homology arms of 469bp (5’ homology arm) and 466bp (3’ homology arm) around the sgRNA1 target site at +9bp from the start of *TBX5* exon 3 was cloned by Cold Fusion (System Biosciences #MC010B) using amplicons from genomic DNA of WTC11 into a construct that was a modification of plasmid pEN114 (Nora et al., 2017). WTC11 cells were electroporated with a cloned nuclease construct containing a guide RNA targeting exon 3, along with the *TBX5* exon3 homology arm-Frt-PGK-EM7-PuroR-bpA-Frt cassette and plated as a serial dilution in mTeSR-1 with Rock inhibitor, as described in (Mandegar et al., 2016). On day 2 and subsequent days, cells were grown in media containing mTeSR-1, Rock inhibitor and puromycin (0.5ug/mL), to select for puromycin-resistant cells. For screening of *TBX5* exon 3 homology-directed repair (HDR) mutations, genomic DNA flanking the targeted sequence was amplified by PCR (For1: ATGGCATCAGGCGTGTCCTATAA, and Rev2: CCCACTTCGTGGAATTTTAGCCA for wildtype, 797 bp, For1: ATGGCATCAGGCGTGTCCTATAA, Rev3: GTTCTTGCAGCTCGGTGAC (Nora et al., 2017) for PuroR, 1631 bp). Positive 5’ arm clones were genotyped by PCR for the 3’ arm (For2: ATTGCATCGCATTGTCTGAG (Nora et al., 2017), Rev4: TTTGACAATCGGGTGGGACC, 829 bp).

For PGP1-derivatives *TBX5^in/+^* or *TBX5^del/del^,* the iPS cell line PGP1 (gift from George Church, available at NIGMS Human Genetic Cell Repository/Coriell #GM23338) (Lee et al., 2009) was electroporated with a cloned nuclease construct containing a guide RNA (sgRNA2) targeting exon 7 of *TBX5*, as described in (Byrne and Church, 2015). For screening of *TBX5* exon 7 NHEJ mutations, the targeted sequence was amplified using PCR primers (For3: GCTTCTTTTGGTTGCCAGAG, Rev5: CATTCTCCCCATTTCCATGT, Seq2: AGAGGCTGCATTTCCATGAT), Illumina compatible-libraries from clones were generated and multiplex-sequenced on a MiSeq for purity of homogeneity of clones for heterozygous or homozygous mutations, as described in (Byrne and Church, 2015).

#### Isolation of homogenous iPS cell clones

Isolation of homogenous colonies for WTC11-derivatives *TBX5^+/+^* (control), *TBX5^in^*^/+^ or *TBX5^in^*^/del^ was performed by modification of methods described previously (Mandegar et al., 2016; Peters et al., 2008). Briefly, single cell suspension of electroporated iPS cells was plated on Matrigel-coated 6 well plates (WP) (BD Bioscience #351146). Once cultures were adherent and recovered to ∼80% confluency, cells were detached by Accutase Cell Detachment Solution (Stemcell Technologies #07920), diluted with 1X DPBS without Ca^2+^/Mg^2+^ and singularized using a P1000 filtered tip, and centrifuged. The cell pellet was resuspended in mTeSR-1, Rock inhibitor and Gentamicin (Life Technologies #15750-060) media, incubated with DAPI (1:1000 from a 1mg/mL stock) for 5 min, centrifuged and resuspended at a concentration of at least 1.0E6 cells/mL in mTeSR-1, Rock inhibitor and Gentamicin media without DAPI. After filtering cells with a 40-micron mesh into FACS tubes, remaining cells (about 120,000 cells per well) were plated onto 6WP for maintenance. Single cells were then sorted for DAPI negativity using a BD FACS AriaII or AriaIII, with a 100-micron nozzle at the lowest flow rate available, into individual wells of a 96WP coated with Matrigel containing media of mTeSR-1, Rock inhibitor and Gentamicin. Upon recovery at 37°C, each well was evaluated one day later for no cells, one cell or more than one cell. All cells were maintained with mTeSR-1, Rock inhibitor and Gentamicin media for at least 5 days, then with mTeSR-1 alone for an additional 5-7 days. Each well at 25% confluency was harvested and re-plated upon singularization with P200 tips in 96WP for more efficient cell growth. When the cell confluency of each well from “single” cells was nearly 100%, then 90% of cells were harvested for genotyping using QuickExtract DNA lysis solution (Epicentre #QE0905T), while 10% of cells were re-plated for the next round of cell selection for wells of interest by FACS sorting again or by serial dilution of cells for manual picking of colonies, as described in (Mandegar et al., 2016; Miyaoka et al., 2014) from apparent “single” cells. Rounds were repeated until every daughter well showed the same genotype, consistent with homogeneity. Genomic DNA from individual wells of interest were amplified using high fidelity *Taq* polymerase, TA-cloned and sequenced to confirm genotype and homogeneity.

Isolation of homogenous colonies for PGP1-derivatives *TBX5^in/+^* or *TBX5^del/del^* was performed as described in (Byrne and Church, 2015). Isolation of homogenous colonies for WTC11-derivative *TBX5^PuR/PuR^* was performed as described in (Mandegar et al., 2016). After sequencing confirmation of respective genotypes, karyotypically-normal cells from each iPS cell line were expanded for subsequent studies.

#### Mice

All mouse protocols were approved by the Institutional Animal Care and Use Committee at UCSF. *Tbx5^del/+^* (Bruneau et al., 2001) and *Tbx5^CreERT2IRES2xFLAG^* (abbreviated here as *Tbx5^CreERT2^*) (Devine et al., 2014b) mice were described previously. *Mef2c^del/+^* mice (Lin et al., 1997) were obtained from Brian Black. *Tbx5^CreERT2/+^* and *Mef2c^del/+^* were maintained in the C57BL6/J background (Jackson Laboratory #664). *Tbx5^fl-bio/fl-bio^* (Waldron et al., 2016) mice were obtained from Frank Conlon. *Mef2a_fl-bio_* and *Mef2c_fl-bio_* (Jackson Laboratory #025983) were described in (Akerberg et al., 2019). *Rosa26BirA* mice were obtained from the Jackson Laboratory (#010920) (Driegen et al., 2005).

### METHOD DETAILS

#### Maintenance of iPS cells and differentiation to cardiomyocytes

All iPS cell lines were transitioned to and maintained on growth factor-reduced basement membrane matrix Matrigel (Corning #356231) in mTeSR-1 medium. For directed cardiomyocyte differentiations, iPS cells were dissociated using Accutase and seeded onto 6WP or 12WP. The culture was allowed to reach 80-90% confluency and induced with the Stemdiff Cardiomyocyte Differentiation Kit (Stemcell Technologies #05010), according to the manufacturer’s instructions. Starting on day 7, differentiations were monitored daily for beating cardiomyocytes and onset of beating was recorded as the day when beating was first observed.

#### Flow Cytometry

iPS-derived cardiomyocytes from WTC11, Control, *TBX5^in/+^* and *TBX5^in/del^* lines were dissociated using Trypsin-EDTA 0.25% on day 15 or day 23 after induction of the differentiation protocol and fixed with 4% methanol-free formaldehyde. Cells were washed with PBS and permeabilized using FACS buffer (0.5% w/v saponin, 4% Fetal Bovine Serum in PBS). For evaluation of differentiation efficiency, cells were stained with a mouse monoclonal antibody for cardiac isoform Ab-1 Troponin at 1:100 dilution (ThermoFisher Scientific #MS-295-P) or the isotype control antibody (ThermoFisher Scientific #14-4714-82). For analyzing levels of Desmin protein, cells were co-stained with the mouse monoclonal antibody for cardiac isoform Ab-1 Troponin at 1:100 dilution and recombinant rabbit anti-Desmin antibody at 1:70 dilution (Abcam #ab32362), or normal rabbit IgG antibody (Millipore Sigma #NI01) for 1 hour at room temperature. After washing with FACS buffer, cells were stained with the following secondary antibodies - goat anti-mouse IgG Alexa 594 at 1:200 dilution (ThermoFisher Scientific #A-11005) and donkey anti-rabbit IgG Alexa 488 at 1:200 dilution (ThermoFisher Scientific #A21206) for 1 hour at room temperature. Cells were then washed with FACS buffer, stained with DAPI for 5 minutes, rinsed, and filtered with a 40-micron mesh. At least 10,000 cells were analyzed using the BD FACSAriaII or AriaIII (BD Bioscience), and results were processed using FlowJo (FlowJo, LLC).

#### Western blotting

iPS-derived cardiomyocytes were harvested on day 15, pelleted and flash frozen. Protein was isolated from supernatant in RIPA buffer with EDTA-free protease and phosphatase inhibitor (ThermoFisher Scientific) after sonication (15 second pulse on, 15 second pulse off, for four pulses). After quantification by BCA assay (ThermoFisher Scientific), 150µg of total protein was loaded per well for each genotype. After running on SDS-PAGE and wet transfer with NuPage Transfer buffer (ThermoFisher Scientific) to a PVDF membrane, the blot was washed in PBST and incubated in primary antibodies of rabbit polyclonal anti-TBX5 at a 1:400 dilution (Sigma #HPA008786) and mouse monoclonal anti-cTNT at 1:1000 dilution (ThermoFisher Scientific #MS-295-P), followed by secondary antibody incubation with donkey anti-rabbit IgG IRDye680 at 1:2000 dilution (Licor #926-68073) and donkey anti-mouse IgG IRDye800 at 1:2000 dilution (Licor #926-32212). The blot was imaged on an Odyssey FC Dual-Mode Imaging system (Licor).

#### Fluorescent *in situ* hybridization

iPS cell-derived cardiomyocytes from WTC11, Control, *TBX5^in/+^*, *TBX5^in/^*^del^ and *TBX5^PuR/PuR^* were dissociated using Trypsin-EDTA 0.25% on day 23 after induction of the differentiation protocol, and 25,000-40,000 cells were plated on to 8-well chambered slides (Ibidi #80826), to obtain a relatively sparse monolayer of cardiomyocytes. Cells were fixed the following day with 10% Neutral Buffered Formalin for 15 minutes at room temperature. Cells were then serially dehydrated in 50%, 70% and 100% ethanol and stored at −20°C until ready to be hybridized. *In situ* hybridization was performed using the RNAscope Multiplex Fluorescent v2 Assay kit (Advanced Cell Diagnostics #323100) with probes for *Hs-TNNT2* (#518991) and *Hs-NPPA* (#531281). Slides were imaged at 10X and 40X magnification on the Keyence BZ-X710 All-in-One Fluorescence Microscope. Non-saturated mean intensity of *NPPA* signal was measured in each *TNNT2+* cell from every group. Unpaired t-tests were used to calculate statistical significance. Brightness and contrast of images in Figure 4N have been adjusted to facilitate viewing of cells.

#### Replating cardiomyocytes for single cell electrophysiology

iPS cell-derived cardiomyocytes (day 15 or older) from WTC11, Control, *TBX5^in/+^*, *TBX5^in/^*^del^ and *TBX5^PuR/PuR^* were gently dissociated in Trypsin-EDTA 0.25% and quenched using StemDiff Maintenance Medium with 10% FBS. Cell suspension was centrifuged at 800 rpm for 5 minutes. The pellet was resuspended in StemDiff Maintenance Medium with Rock inhibitor at a 1:1000 dilution. Cardiomyocytes were counted, and 25,000-35,000 cells were plated on to growth factor-reduced Matrigel-coated 15mm round glass coverslips (Warner Instruments #64-0703) to obtain a sparse distribution. Cardiomyocytes were then maintained on coverslips in StemDiff Maintenance Medium.

#### Patch Clamp Electrophysiology

Patch clamp recordings were made on single iPSC-derived cardiomyocytes using the perforated-patch configuration. Experiments were performed at 30°C under continuous perfusion of warmed Tyrode’s solution containing (in mM): 140 NaCl, 5.4 KCl, 1 CaCl_2_, 1 MgCl_2_, 10 glucose, and 10 HEPES, with the pH adjusted to 7.4 with NaOH. Recordings were conducted using borosilicate glass pipettes (Sutter Instruments) with typical resistances of 2 to 4MW. The pipette solution consisted of (in mM): 150 KCl, 5 NaCl, 5 MgATP, 10 HEPES, 5 EGTA, 2 CaCl_2_, and 240 mg/mL amphotericin B, with the pH adjusted to 7.2 with KOH. Spontaneous action potentials were acquired in a zero-current current clamp configuration using an Axopatch 200B amplifier and pClamp 10 software (Axon Instruments). Data was digitized at 20 kHz and filtered at 1kHz. Action potential parameters from each cell were derived using Clampfit 10 software (Axon Instruments).

#### Calcium imaging

iPSC-derived cardiomyocytes on glass coverslips were loaded with Ca^2+^ indicator dye Fluo-4 AM (Thermo Fisher Scientific #F14201) to record Ca^2+^ flux, as previously described (Spencer et al., 2014). Measurements were made on spontaneously firing single or small clusters of iPSC-derived cardiomyocytes using a 10X objective on a Zeiss Axio Observer Z1 inverted microscope. For experiments, cells were placed in Tyrode’s solution containing 1.8 mM Ca^2+^ within a 37°C heated stage-top imaging chamber (Okolab). Images were acquired at 100 fps using an ORCA-Flash 4.0 camera (Hamamatsu, Bridgewater, NJ). Data was processed using ZEN (Zeiss) or Image J software (http://rsbweb.nih.gov/ij/) and analyzed using custom in-house software (Hookway et al., 2019).

#### Immunostaining of cardiomyocytes

iPSC-derived cardiomyocytes from WTC11, Control, TBX5^in/+^ and *TBX5^in/de^*^l^ were replated on coverslips placed in 12-well plates on day 23, as described above for replating for electrophysiology. Cells were fixed in 4% formaldehyde for 20 minutes at room temperature, followed by washes in PBS. Cells were then treated with a blocking buffer containing 5% goat serum and 0.1% Triton X-100 in PBS for 1 hour at room temperature. A mouse monoclonal antibody for cardiac isoform Ab-1 Troponin (ThermoFisher Scientific #MS-295-P) was added to the coverslip-containing wells at a 1:100 dilution in blocking buffer and incubated on a rocker for 2 hours at room temperature. Following washes with 0.1% Triton X-100 in PBS, coverslips were treated with a donkey anti-rabbit IgG Alexa 488 antibody (ThermoFisher Scientific #A21206) at a 1:200 dilution for 2 hours at room temperature. Coverslips were then washed with 0.1% Triton X-100 in PBS and stained with DAPI at a 1:1000 dilution for 2 minutes. Coverslips were washed and stored in PBS at 4C. Images were acquired on a Zeiss LSM 880 with Airyscan and processed by ImageJ (Abràmoff et al., 2004).

#### Cell harvesting for single cell RNA sequencing

Cells from day 6, day 11 or day 23 of the differentiation protocol were collected from 3 independent differentiations. Wells for dissociation were chosen based on typical differentiated morphology on day 6 or robust beating on day 11 and day 23. Cells were singularized with Trypsin-EDTA 0.25%. After quenching, the single cell suspension was centrifuged at 800 rpm for 5 minutes. The pellet was resuspended in 1X PBS with 0.04% w/v Ultrapure BSA (MCLAB #UBSA-500) and counted. A 30µL cell suspension containing 10,000 cells was used to generate single cell droplet libraries with the Chromium Single Cell 3′ GEM, Library & Gel Bead Kit v2 according to manufacturer’s instructions (10X Genomics). After KAPA qPCR quantification, a shallow sequencing run was performed on a NextSeq 500 (Illumina) prior to deep sequencing on a NextSeq 500, HiSeq 4000, or NovaSeq (Illumina) for a read depth of >100 million reads per cell.

#### Data processing using Cellranger

All datasets were processed using Cellranger 2.0.2. FASTQ files were generated using the mkfastq function. Reads were aligned to hg19 reference (version 1.2.0). Cellranger aggr was used to aggregate multiple GEM libraries.

#### Seurat analysis

Outputs from the Cellranger pipeline were analyzed using the Seurat package (version 2.3.4 or 3.1.4) (Butler et al., 2018; Satija et al., 2015; Stuart et al., 2019) in R (version 3.5.1) [R Core Team (2018). R: A language and environment for statistical computing. R Foundation for Statistical Computing, Vienna, Austria. URL https://www.R-project.org/]. Datasets from day 6, day 11 or day 23 experiments were analyzed as separate Seurat objects. Seurat objects for day 6 or day 11 were generated using Seurat v2. Seurat objects for day 23 datasets with multiple biological replicates were generated using Seurat v3, unless otherwise noted.

Quality control steps were performed to remove dead cells or doublets, and cells with a UMI count between 10,000 to 80,000 were retained. After normalizing the data, sources of unwanted variation, such as differences in the number of UMI, number of genes, percentage of mitochondrial reads and differences between G2M and S phase scores were regressed using the ScaleData function. Next, principal component analysis (PCA) was performed using the most highly variable genes. Cells were then clustered based on the top 25-30 principal components and visualized using a dimensionality reduction method called Uniform Manifold Approximation and Projection (UMAP) (Becht et al., 2018). The resolution parameter was set, so that cluster boundaries largely separated the likely major cell types.

Two technical replicates at day 6 and day 11 for WTC11-derived cells (WTC11, control, *TBX5^in/+^*, *TBX5^in/del^*) were evaluated. For control at day 23, two technical replicates were evaluated. For WTC11, *TBX5^in/+^* or *TBX5^in/del^* at day 23, two technical replicates from biological replicate 1 and one sample from biological replicate 2 were evaluated. For PGP1-derived cells (PGP1, *TBX5^in/+^* and *TBX5^del/del^*), one sample for each genotype was evaluated.

Major cell type categories were defined by their expression of select enriched genes in a given cluster--pluripotent cells (*POU5F1*) cardiomyocytes (*TNNT2*), dividing cardiomyocytes (*CENPF*^+^/*TNNT2^+^*), ventricular cardiomyocytes (*TNNT2^+^/IRX4^+^*), fibroblasts (*COL1A1*), epicardial cells (*WT1*^+^/*TBX18*^+^), neural crest-derived cells (*MAFB^+^*, *MSX1^+^*), endoderm (*TTR* alone or *TTR*^+^/*AFP^+^*) and endothelial cells (*PLVAP*). Clusters of cells not defined by any of these markers were labeled as “Undetermined”. The numbers of cells in each major cell type category in each genotype were then calculated. Sunburst plot was generated in Excel using the percentage of cells in each cell type category per genotype. We used FindAllMarkers to generate a list of top marker genes for each cluster and highlighted selected genes in a Manhattan plot to display potential diversity of subtypes among these major cell types.

#### Integration and Visualization of Datasets from Multiple Samples

For the day 23 WTC11-derived cell line (biological replicate 1 and 2) analysis, we ran CellRanger to normalize sequencing depth variation between individual libraries. We then ran Seurat v3.1.4’s ‘Integration and Label Transfer-SCTransform’ workflow to resolve effects from experimental instances that are driven by cell-cell technical variations, including sequencing depth (Hafemeister and Satija, 2019; Stuart et al., 2019). Cells with lower than 10,000 UMIs and concurrently higher percentage of mitochondrial reads were removed. Potential doublets with higher than 75,000 UMIs were also removed. The dataset was then split into two Seurat objects using the biological replicate status. We ran CellCycleScoring(default) and SCTransform(vars.to.regress=c(“S.Score”, “G2M.Score”)) to regress out cell cycle variations.

The remaining steps followed the ‘Integration and Label Transfer-SCTransform’ workflow. Briefly, these steps include finding 2,000 highly variable genes to create anchors that represent biologically common cells connected from opposing batches. After integration, Seurat set the active assay to ‘integrated’ for downstream data visualization analysis. UMAPs were created by running RunPCA(default) and RunUMAP(default).

We also evaluated genetic backgrounds from two iPSC lines. The WTC11-derived cell lines were considered genetic background 1, which included biological replicate 1 and 2. PGP1-derived cell lines were considered genetic background 2. We followed the same CellRanger aggregate and qc filtering. However, we used the genetic background status to make three Seurat objects and no variables were regressed when running the ‘Integration and Label Transfer-SCTransform’ workflow. UMAPs were created by running RunICA(default) and RunUMAP(reduction=”ica”, dims=1:40, min.dist=0.4, spread=0.9, repulsion.strength=6).

For day 23 cardiomyocyte datasets, *TNNT2^+^* clusters were defined as containing a majority of cells expressing *TNNT2* on a feature plot and extracted using the subset function and re-clustered. Subsequently, the resolution parameter was set to partition clusters enriched for a particular genotype. A phylogenetic tree was generated by relating the “average” cell from each cluster in PC space, using the BuildClusterTree function. Differential gene expression tests were run between closely related clusters, using the FindMarkers function with min.pct set to 0.1 and logfc.threshold set to 0.25. Selected differentially expressed genes with an adjusted p-value less than 0.05 from the Wilcoxon Rank Sum test were then displayed using the Dotplot function. As Seurat log normalizes gene expression counts and scales values for each gene (mean is 0, std dev of +/-1), dot plots and heatmaps are based on scaled expression values.

#### Cell Type Classifier by Machine Learning

We applied machine learning to predict corresponding *in vivo* cell types in our WTC11-derived samples. A sklearn multiclass logistic regression model, using a one-vs-rest scheme and the cross-entropy loss cost function (Pedregosa et al., 2011), was trained on the *in vivo* scRNA-seq dataset published by (Asp et al., 2019). The training data contained eleven cardiac cell type classes (i.e. Fibroblast-like, atrial cardiomyocyte (aCM)-like, ventricular cardiomyocyte (vCM)-like, Cardiac neural crest-like, Sub-epicardial-like, Capillary endothelium/pericytes/adventitia-like, Smooth muscle/fibroblast-like, and Erythrocyte-like). The test data was the day 23 integrated WTC11-biological replicates.

We ran SCTransform(default) independently on the training and test data to remove sequencing depth bias, while preserving biological heterogeneity. To train our classifier on cell-type specific signals from both datasets, we used SCTransform Pearson residuals as the feature space for 1,538 genes. The genes were selected by taking the intersection of the top 3,000 highly variable genes (HVGs) from the training and test datasets (Table S1).

We evaluated the cell type classifier using sklearn’s stratified 10-fold cross validation method; StratifiedKFold(n_splits=10, random_state=42). Each fold preserves the percentage of *in vivo* cell types. Thus, recapitulating true *in vivo* cardiac cell type composition in our training evaluation. For each fold of the cross validation, we used a sklearn logistic regression model to fit and predict on the fold’s training and test set; LogisticRegression(penalty=’l2’, solver=’lbfgs’, random_state=42). The cross validation model’s average performance measurements were: accuracy (96.9%), precision (97.8%), recall (97.4%), and f1 score (97.1%) (Figure S2C). Due to the strong cross validation performance, we trained our deployment model on the full *in vivo* dataset to increase cell type generalisability. The trained multinomial classifier was then deployed on our WTC11-derived samples.

#### Congenital Heart Disease-Associated or Electrophysiology-Related Gene Lists and Cell Type Expression

A list of 375 CHD candidate genes, including inherited, *de novo*, syndromic or non-syndromic CHD genes of interest, was manually curated from literature (Homsy et al., 2015; Jin et al., 2017; Lalani and Belmont, 2014; McCulley and Black, 2012; Prendiville et al., 2014; Priest et al., 2016; Sifrim et al., 2016; Zaidi et al., 2013). A list of 76 EP genes were manually curated. A list of cardiac development-related factors is from (Duan et al., 2019). Lists can be found in Table S3.

#### Cell trajectories and pseudotime analysis

Pseudotime analysis was performed using the URD package (version 1.0.2) (Farrell et al., 2018b). A single Seurat object (from Seurat v2), consisting of combined data from two technical replicates of three timepoints and four genotypes, was processed as described in the previous section, and then converted to an URD object using the seuratToURD function. Cell-to-cell transition probabilities were constructed by setting the number of nearest neighbors (knn) to 211 and sigma to 8. Pseudotime was then calculated by running 80 flood simulations with *POU5F1^+^* clusters as the ‘root’ cells. Next, all day 23 clusters were set as ‘tip’ cells and biased random walks were simulated from each tip to build an URD tree.

We identified URD monotonic genes, which are genes that neither deviate from an increase or decrease in expression with pseudotime. Spearman rank correlation (Python v3.7.3, and libraries Pandas 0.25.0, Numpy 1.17.1, and SciPy 1.3.1) was used to find significant monotonic genes (p-value < 0.05). To determine if these monotonic relationships differ between WT and *TBX5^in^*^/*del*^ paths to cardiomyocytes, we used a Fisher z-transformation to test the null hypothesis that there is no significant difference in correlation (Fisher, 1921). To illustrate these results, we use heatmaps for genes with a |rho|≥0.4 to pseudotime and Z-score≥15 as a difference between WT and *TBX5^in/del^* paths.

To identify differential expressed genes in inferred cardiac precursors (intermediate branches in the URD tree) that are affected by *TBX5* loss, cell barcodes from each precursor segment (wildtype/control/*TBX5^in/+^* path vs. *TBX5^in/del^* path) were extracted from the URD object and assigned new identities in the corresponding Seurat object. Differential gene test was then performed between the two segments using Wilcoxon Rank Sum test with min.pct set to 0.1 and logfc.threshold set to 0.25. Selected genes with an adjusted p-value less than 0.05 were plotted on the URD tree to visualize their expression.

To compare the trident (*TNNT2*^+^ distal branch for WTC11, control and *TBX5^in/+^*) and fork (*TNNT2*^+^ distal branch for *TBX5^in/del^*) during pseudotime, we subdivided the pseudotime from the common branchpoint to the tips of the trident and fork into twenty uniform windows. Within each window, we then calculated the *t* test, difference of means, and fold change between the trident and fork for all genes. We filtered the statistics by gene-window combinations with adjusted p-value<0.05 after Bonferroni-Holm multiple testing correction. Then, we hierarchically clustered the genes on *t* test p-values and plotted statistics using the R pheatmap library.

#### Cell browser implementation

The cell browser at cells.ucsc.edu was developed by Maximilian Haeussler. We created a cell browser session that allows the user to interrogate the spatial distribution of metadata and expression across data, in multiple reduced dimensionality spaces including the URD trajectory. Using a Scanpy python pipeline, we generated PCA, tSNE, UMAP, PAGA, and drl transforms. We also imported the URD trajectory mapping and WGCNA transform from their respective packages. We ran the scoreCT algorithm to assign cell types to cell clusters using a marker gene set.

#### Gene regulatory network analysis

bigSCale2 (https://github.com/iaconogi/bigSCale2) (Iacono et al., 2019; 2018) was used with default parameters to infer gene regulatory networks and “correlomes” from single cell RNA-seq expression data for *TNNT2*^+^ cells. Expression counts and gene names were used as input from two technical replicates at each time point and *TBX5* genotype. Details of each dataset can be found in Table S6. To evaluate significant changes in pagerank or degree centrality, we computed all pairwise differential differences in pagerank or degree between baseline (wildtype and control) vs. *TBX5* mutants (*TBX5^in/+^* or *TBX5^in/del^*) (12 total differences, from 2 *TBX5* mutants * 2 baselines * 3 stages) and used these values to determine the top 5% upper change cutoff from 8,704 genes of all networks. Classification of Pearson correlations were empirically chosen at >0.5 for correlation and <-0.05 for anti-correlation.

#### ChIP-seq

Combined peaks of human TBX5 or GATA4 ChIP-seq from hiPSC-derived cardiomyocytes were used (Ang et al., 2016). bioChIP-seq of mouse TBX5, MEF2c and MEF2a from E12.5 hearts were from (Akerberg et al., 2019). Single replicates of TF bioChIP peaks, which were IDR normalized (IDR_THRESHOLD=0.05 between each set of replicates), were defined as the summit of the peak with the strongest ChIP signal ± 100bp of the individual replicate with the greatest peak intensity. Mouse H3K27ac ChIP-seq at E12.5 of embryonic cardiac ventricles was from (He et al., 2014).

### QUANTIFICATION AND STATISTICAL ANALYSIS

#### Scoring of sarcomeric disarray

Myofibrillar arrangement in cardiomyocytes was manually scored on a scale of 1-5, similar to (Judge et al., 2017). A score of 1 represents cells with intact myofibrils in a parallel arrangement. A score of 2 represents cells that have intact myofibrils, but many are not parallel. Scores of 3 and 4 include cells with increasing degrees of myofibrillar fragmentation or aggregation. A score of 5 represents cells without visible myofibrils. No cells were apparent among our samples with a score of 5. Violin plots were generated in Prism (GraphPad) to show distribution of scored cells from each group. Fisher’s exact test was used to determine statistical significance.

#### Graphing and statistics for electrophysiology

For electrophysiology and calcium imaging experiments, graphs were generated using Prism 8.2.0 (GraphPad Software). Significance between parental and experimental groups was determined with a custom R-script using unpaired two-sided Welch’s *t* tests with Holm-Sidak correction for multiple comparisons (Holm1979). Adjusted p-value<0.05 was considered statistically significant.

#### Statistical analyses for correlations

We evaluated the pairwise association among 38 variables, including all human genes, TBX5-dysregulated genes in human cardiomyocytes from day 23, CHD genes, EP genes, TBX5 or GATA4 binding (Ang et al., 2016), and genome-wide association (GWAS) genes for CHDs or arrhythmias (Cordell et al., 2013a; 2013b; Ellinor et al., 2012; Hoed et al., 2013; Hu et al., 2013; Pfeufer et al., 2010a; Smith et al., 2011). Reported GWAS genes from https://www.ebi.ac.uk/gwas/ for the terms congenital heart disease, congenital heart malformation, congenital left sided heart lesions, conotruncal heart defect and aortic coarctation were used to define congenital heart disease-related (CHD) GWAS genes. Reported genes from terms such as cardiac arrhythmia, supraventricular ectopy, ventricular ectopy, premature cardiac contractions, atrial fibrillation, sudden cardiac arrest and ventricular fibrillation were considered as arrhythmia-related (EP-GWAS) genes. Two nearest genes within 100kb, by using GREAT (great.stanford.edu) (McLean et al., 2010), of TBX5 or GATA4 binding sites or of reported genes from each group of GWAS, were considered for the analysis. The natural logarithm odds of genes associating with each one of these variables versus the odds of genes associating with every other variable were estimated using generalized linear models with family=“binomial” setting in R. The resulting significance of these natural log odds ratios were adjusted for multiple testing by the Benjamini-Hochberg method (Benjamini and Hochberg, 1995). Significance was determined using an FDR threshold of 0.05 or less.

Additional correlations were evaluated between 26 variables, including human genes, human TBX5-dysregulated genes from day 23 cardiomyocytes, and TBX5, MEF2c or MEF2a binding sites from E12.5 mouse heart tissue (Akerberg et al., 2019). Human gene symbols were converted to mouse gene symbols, using the getLDS() function from the biomaRt package (https://www.r-bloggers.com/converting-mouse-to-human-gene-names-with-biomart-package/). Two nearest genes within 100kb of TBX5, MEF2c or MEF2a binding sites were considered for the analysis.

For assessment of associations between binding locations of TBX5, MEF2c and MEF2a transcription factors with genes dysregulated by TBX5, analyses were performed corresponding to binding regions of each of the three TFs. First, binding regions of each TF was evaluated for association with genes, defined by the nearest two genes within 100kb. Using the list of human TBX5-dysregulated genes, binding regions of each TF associated with a TBX5-dysregulated gene was determined. Identified binding regions of each TF that overlapped with at least 50% of the binding regions of each of the other two TFs was determined, using bedops --element-of - 50%. This approach defined three variables, including every binding region of the TF, if associated with a TBX5-dysregulated gene, or if it overlaps by at least 50% with the binding region of the other TFs, that were used for logistic regression in R. The resulting changes in odds are represented as natural logarithm odds ratios. Multiple testing correction was performed using the multtest package in R. All estimates are based on analyses for human TBX5-dysregulated genes.

### DATA AND SOFTWARE AVAILABILITY

scRNA-seq datasets have been deposited at NCBI GEO, under accession GSE137876. R and python scripts will be available upon publication.

---

**Figure S1.**
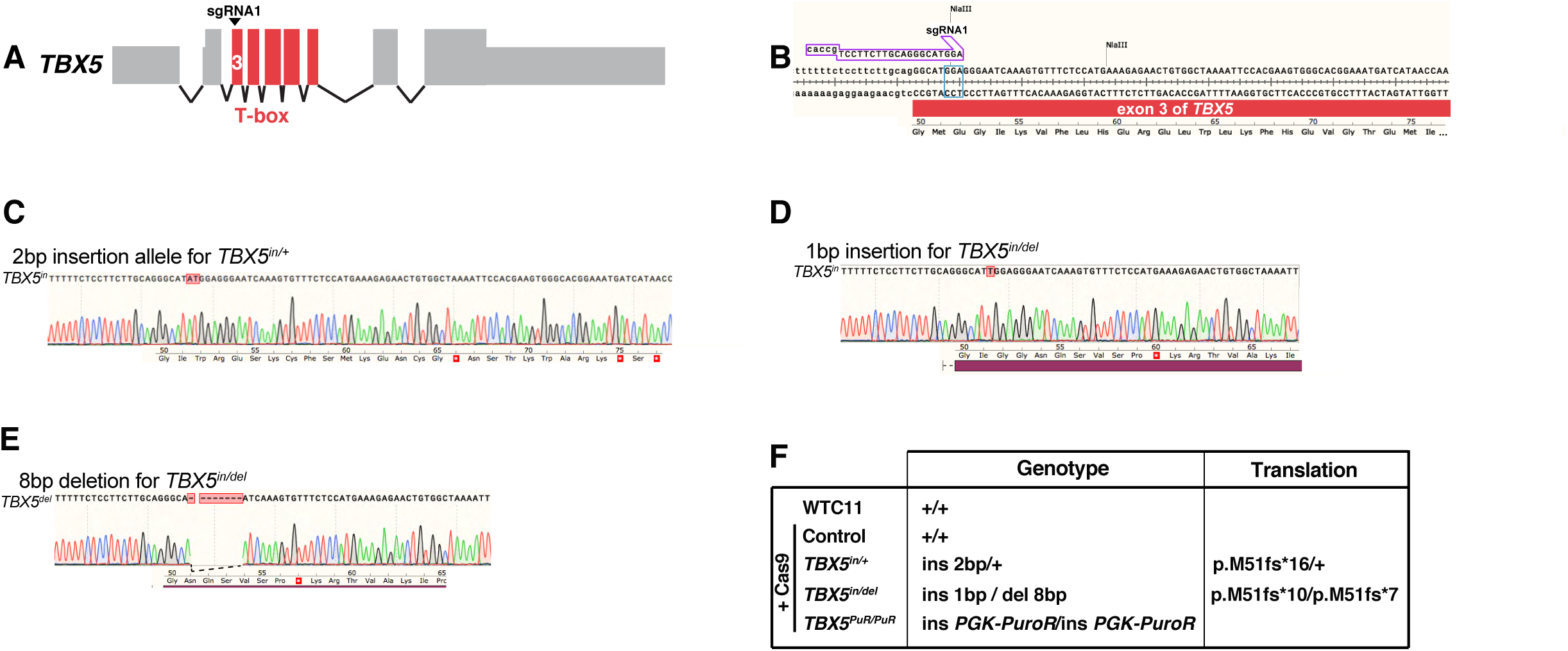
Genome editing of *TBX5* in human induced pluripotent stem cells. (A) Diagram of the human *TBX5* gene. Exons encoding the T-box domain of TBX5 are indicated in red. sgRNA1 was used to target exon 3 of *TBX5* by a CRISPR/Cas9 nuclease. (B) Sequence of the exon 3 of *TBX5* is shown, along with the sgRNA1 location. The PAM site is boxed in blue. Loss of the NlaIII site at the PAM site was used in initial screening for mutant iPS cell clones by PCR. The encoded wildtype protein sequence includes the start of the T-box domain. (C) Sequence and chromatogram for the 2bp insertion of the mutant allele for *TBX5^in/+^* predicts a premature truncation, as indicated by a stop codon (white asterisk in red box) in the frame-shifted protein sequence. (D, E) Sequence and chromatogram for the 1 bp insertion, or 8 bp deletion, respectively, of the mutant allele for *TBX5^in/del^*, along with corresponding protein sequences, are shown. (F) Table shows genotypes of WTC11-derived iPS cell lines that were targeted for *TBX5* at exon 3. Predicted translation for each *TBX5* genotype is indicated.

**Figure S2.**
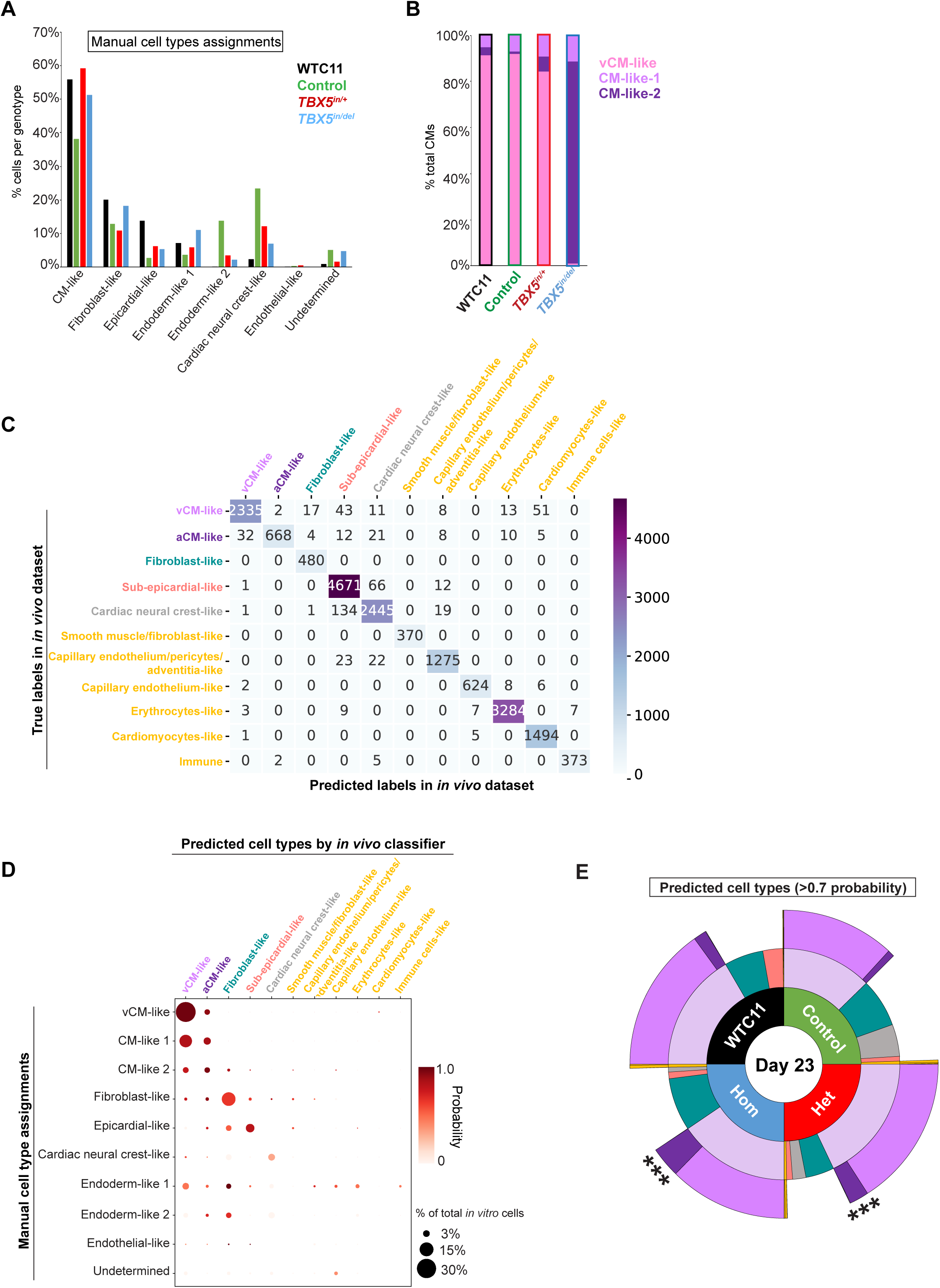
Diversity of iPSC-derived cell types by *TBX5* genotype. (A) Manual assignment of iPSC-derived cell types by *TBX5* genotypes at day 23. (B) Distribution of iPSC-derived cardiomyocyte classification by *TBX5* genotype at day 23. (C) A confusion matrix compares test vs. predicted cell type labels for human fetal cardiac cells (Asp et al., 2019). (D) A confusion matrix compares cell type assignments of iPSC-derived cells at day 23 by manual annotation and *in vivo* classifier prediction. Color of each dot represents a prediction probability of the *in vivo* cell type classifier, while the dot size displays the percentage of the total iPSC-derived cells at day 23. (E) Distribution for predicted cell types of >0.7 prediction probability is shown by *TBX5* genotype at day 23 in a sunburst plot. *** p<0.0001 by Fisher’s exact test.

**Figure S3.**
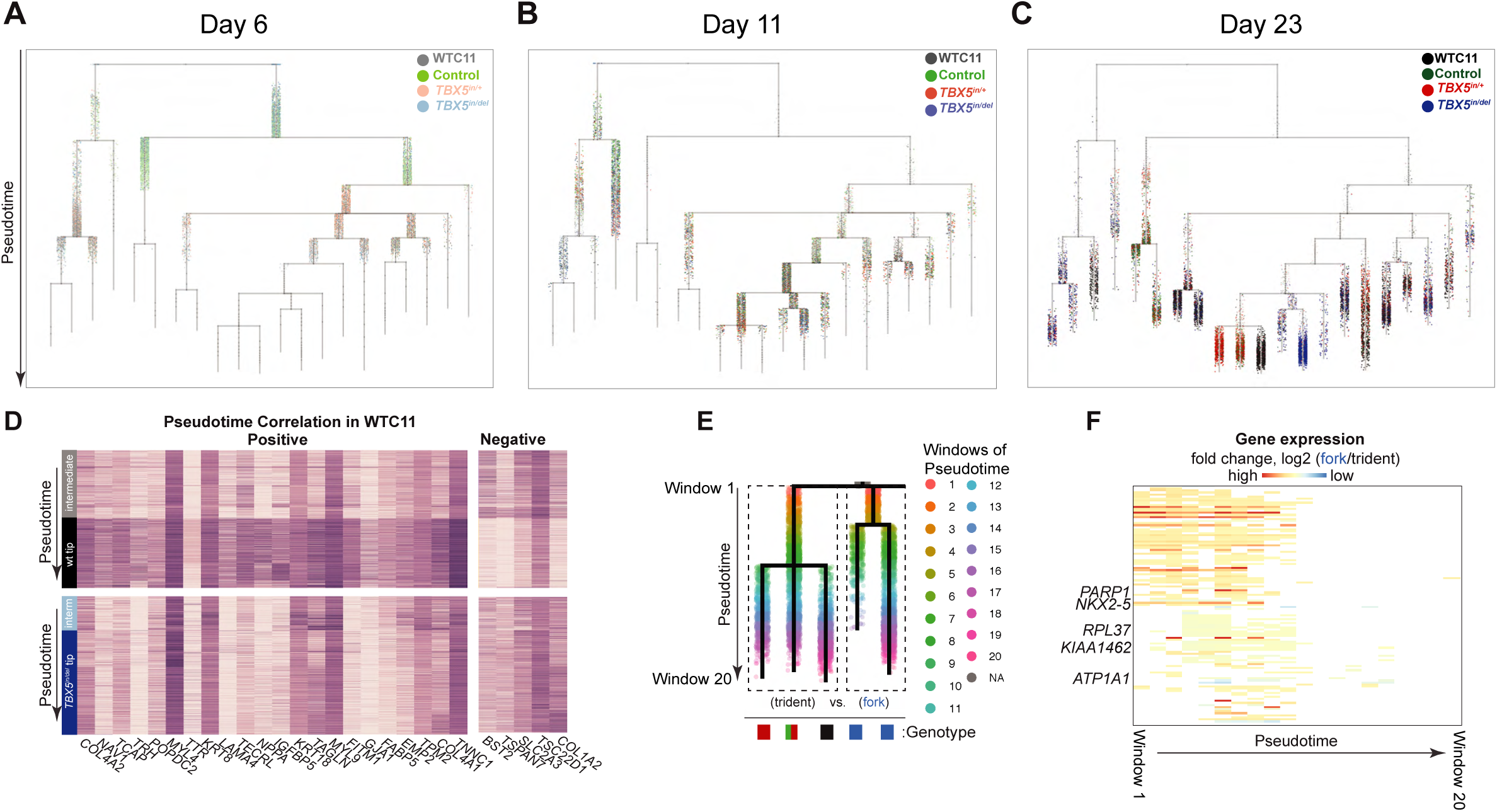
Pseudotime analysis of TBX5-dependent cardiomyocyte differentiation. (A-C) Cells from all *TBX5* genotypes at day 6, 11 or 23 are shown by harvested time point on an aggregate pseudotime dendrogram using URD trajectory inferences. (D) Heatmaps show expression for each gene that displays a positive or negative correlation with pseudotime (|rho|≥0.4 and Z-score≥15 by difference in rho) in the WT path (above) and is altered in the *TBX5^in^*^/*del*^ path (below). (E) Paths for WT/*TBX5^in^*^/*+*^(trident) or *TBX5^in^*^/*del*^ (fork) to cardiomyocytes were divided into windows (1-20) along pseudotime for comparison. (F) Heatmap shows fold change for genes in a cluster that includes *NKX2-5,* which was significantly different after correction (adj p-value<0.05 by Bonferroni-Holm test) in windows 2 through 8 between the deduced WT/control/ *TBX5^in^*^/*+*^ and *TBX5^in^*^/*del*^ paths, along with genes of a similar pattern, including *PARP1, RPL37, KIAA1462,* and *ATP1A1* (adj p-value<0.05 by Bonferroni-Holm test).

**Figure S4.**
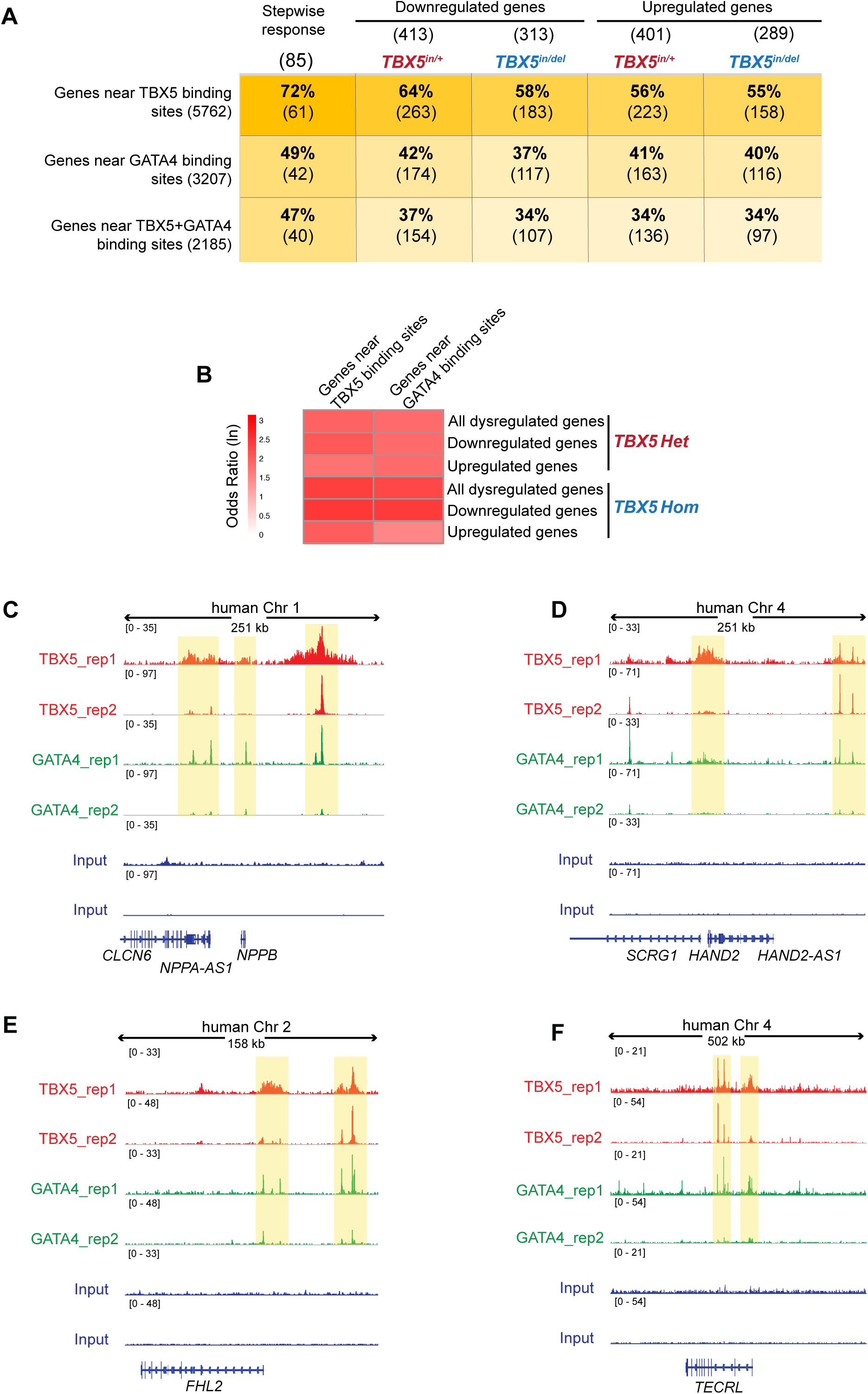
TBX5 and GATA4 occupancy near human TBX5-dependent genes. (A) Table shows number of TBX5-dependent genes near TBX5, GATA4 or TBX5 and GATA4 occupancy (Ang et al., 2016). (B) Heatmap displays significant correlations (FDR<0.05) of TBX5 or GATA4 occupancy near human TBX5-dependent gene sets (Table S5). Odds ratios for co-occupancy of TBX5 and GATA4 at TBX5-dependent genes are in Table S6. (B-E) Browser tracks of TBX5 and GATA4 occupancy from iPSC-derived cardiomyocytes are shown for loci of TBX5-dependent genes *NPPA/NPPB*, *HAND2, FHL2* and *TECRL*.

**Figure S5.**
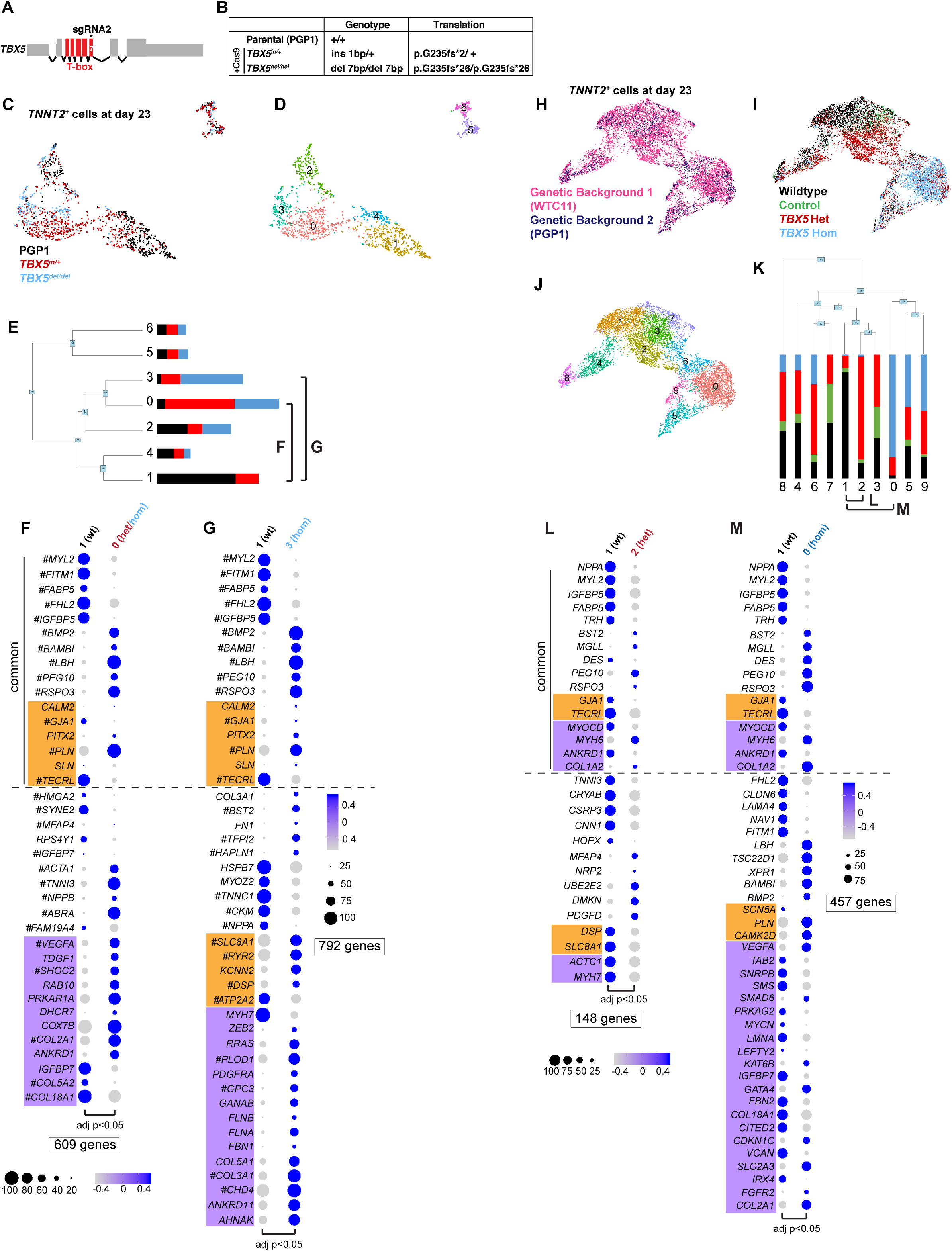
Assessment of two genetic backgrounds for TBX5 dose-sensitive gene expression. (A) Diagram of the human *TBX5* gene is shown, with exons in red. The guide sgRNA2 was used to target exon 7, which encodes a portion of the T-box domain, of *TBX5* in PGP1 iPS cells. (B) A table specifies each *TBX5* mutation and the predicted translation of TBX5 for PGP1-derived *TBX5^in/+^* or *TBX5^del/del^* cells. (C, D) UMAPs of *TNNT2^+^* cells at day 23 by *TBX5* genotype (C) or cluster identity (D). (E) A phylogenetic tree shows the relatedness of the ‘average’ cell in each cluster using PC space. The percentage of cells within a cluster from each *TBX5* genotype are colored. Related clusters between different *TBX5* genotypes were compared for differential gene expression. (F, G) Dot plots show top differentially expressed genes in (F) *TBX5^in/+^*- or (G) *TBX5^del/del^*-enriched clusters. Significance was determined by Wilcoxon Rank Sum test (adj p-value<0.05) (Table S2). (H-J) *TNNT2*^+^ cells are displayed in a UMAP, by genetic background (WTC11 or PGP1-derived cells) (H), by *TBX5* genotype (I), or by Louvain clustering (J). (K) A phylogenetic tree shows the relatedness of the ‘average’ cell in each cluster using PC space. The proportion of cells in each cluster are colored by *TBX5* genotype. Related clusters between different *TBX5* genotypes were selected for differential gene tests. (L, M) Dot plots show top differentially expressed genes in (L) *TBX5^in/+^*- or (M) *TBX5^del/del^*-enriched clusters. Top five upregulated or downregulated differentially expressed genes, along with EP and CHD genes, were common between comparisons. Significance was determined by Wilcoxon Rank Sum test (adj p-value<0.05) (Table S2, S3).

**Figure S6.**
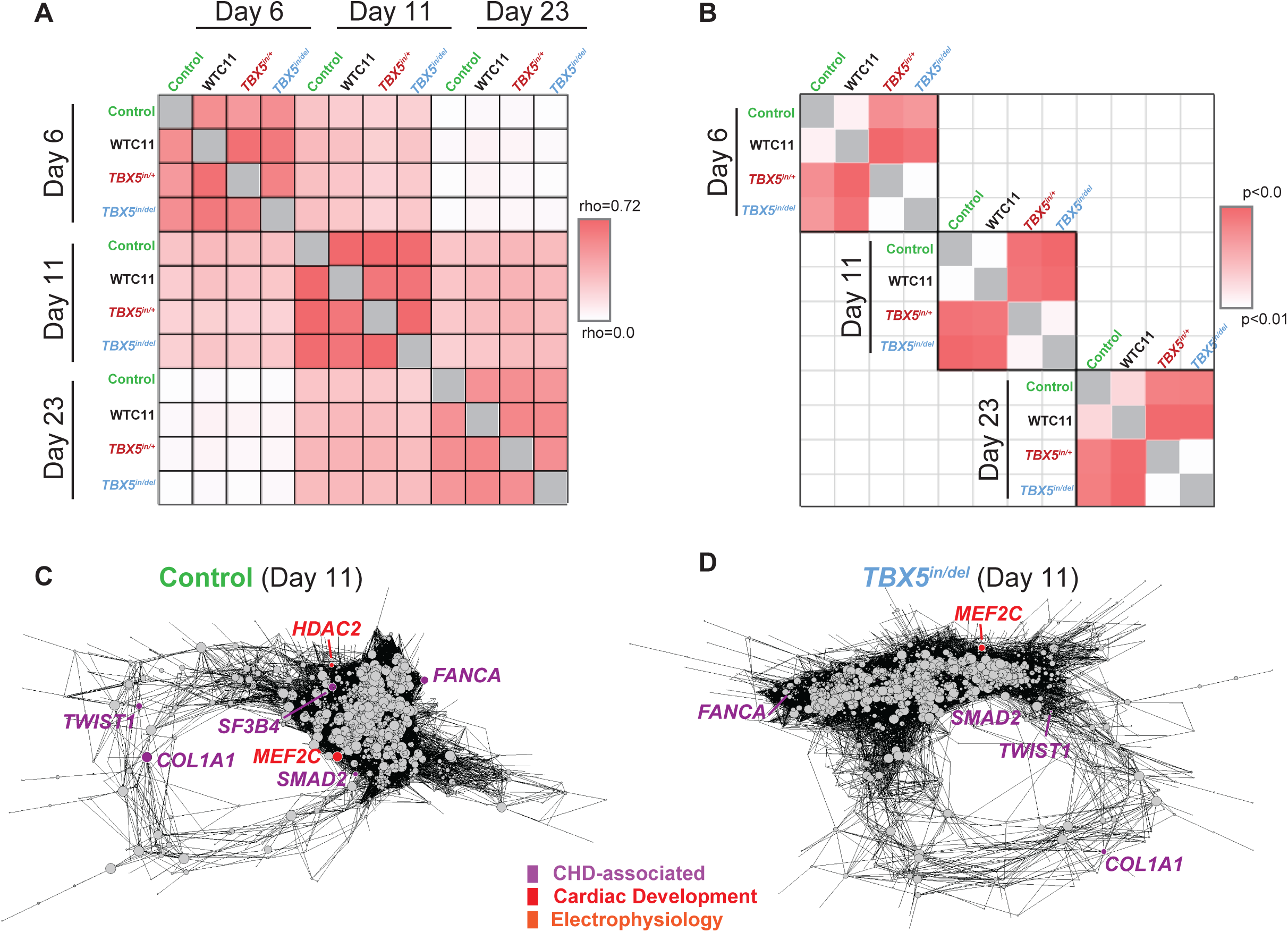
Analysis of TBX5 dosage-sensitive gene regulatory networks. (A) Correlation plot (Pearson correlations of pagerank centralities) of *TNNT2^+^* networks by *TBX5* genotypes and time points are shown. Note that networks display highest similarity (red) within a time point. An inter-stage dissimilarity (white) grows proportionally to the time difference (i.e. day 23 is less similar to day 6 than to day 11). Therefore, comparisons for genotype differences were made within differentiation stages. (C) Network similarity among *TBX5* genotypes within each time point is shown (Wilcoxon Rank Sum test of pagerank centralities for nodes from selected time point comparisons). (C, D) Network diagrams of *TNNT2^+^* cells at day 11 for control (C) or *TBX5^in/del^* (D) are shown.

## Supplementary Tables

Table S1. Classifier gene features and weights for each cell type prediction.

Table S2. Lists of differential genes from comparisons between *TNNT2*^+^ clusters, at day 23 by biological replicate (Figure 4) or genetic background at day 23 (Figure S5).

Table S3. Curated gene lists, which are used in this study, include electrophysiology (EP) genes, human congenital heart disease (CHD) genes, mouse CHD genes, cardiac development genes, CHD-associated GWAS genes and EP-related GWAS genes.

Table S4. Correlation of human TBX5-dependent genes near TBX5 or GATA4 occupancy, congenital heart disease (CHD)-associated GWAS, electrophysiology (EP)-related GWAS, CHD genes, or EP genes. Odds ratios are displayed as natural logarithms. Statistical significance was determined by Benjamini-Hochberg multiple testing.

Table S5. Odds ratio as natural logarithm for transcription factor (TF) binding of TBX5 or GATA4 within 1kb of the other TF, near human TBX5-dependent genes. Statistical significance was determined by Benjamini-Hochberg multiple testing.

Table S6. TBX5-sensitive gene regulatory network analyses, by pagerank or degree, or by correlation with *TBX5* or *MEF2C* expression.

Table S7. Correlation of mouse orthologs of human TBX5-dependent genes near TBX5, MEF2c or MEF2a occupancy, congenital heart disease (CHD)-associated GWAS, electrophysiology (EP)-related GWAS, CHD genes, or EP genes. Odds ratios as natural logarithms are displayed. Statistical significance was determined by Benjamini-Hochberg multiple testing.

Table S8. Odds ratio as natural logarithms for transcription factor (TF) binding of TBX5, MEF2c, or MEF2a within 1kb of the other TFs in the trio, near mouse orthologs of human TBX5-dependent genes. Statistical significance was determined by Benjamini-Hochberg multiple testing.

